# Dynamic Retinal Pathology in Glaucoma Progression Revealed by High-Resolution Functional Imaging *in Vivo*

**DOI:** 10.1101/2025.07.22.666227

**Authors:** Yiming Fu, Pham Binh Minh, Sicong He, Yingzhu He, Zhongya Qin, Ting Xie, Jianan Qu

## Abstract

Glaucoma, a complex and multifactorial neurodegenerative disease, is commonly associated with elevated intraocular pressure and primarily characterized by the progressive loss of retinal ganglion cells (RGCs) and their axons. Despite its prevalence, our understanding and treatment of this disease remain challenging due to the intricate interplay of various pathophysiological factors and the limited capability for *in vivo* functional study of the disease. In this work, we investigated the dynamic retinal pathology of glaucoma from onset to late stages through longitudinal *in vivo* high-resolution imaging in a silicone oil-induced ocular hypertension glaucoma mouse model. We developed an optimized adaptive optics two-photon excitation fluorescence microscopy (AO-TPEFM) technique for both morphological and functional assessments of pathological changes in the retina across three distinct glaucoma phenotypes with varying progression rates. Our AO-TPEFM technology visualized the complete process of functional and structural changes in the three most important retinal components related to glaucoma: microvascular vessels, microglia, and RGCs during disease progression. Notably, our functional imaging revealed pathological alterations in microvascular circulation, microglia state, and RGC functionalities at a very early stage of disease development when retinal morphology still appeared normal, providing critical insights into glaucoma pathogenesis. This research also demonstrates that AO-TPEFM is a powerful tool for the *in vivo* study of general retinal diseases.

## Introduction

Glaucoma encompasses a group of ocular disorders that constitute the most common cause of irreversible blindness, affecting nearly 95 million people worldwide^1–3^. Elevated intraocular pressure (IOP) represents the primary risk factor for glaucoma^4^ and the disease is characterized by the degeneration of retinal ganglion cells (RGCs), the only projection neurons in the retina to brain^5–7^. Unfortunately, glaucoma is often diagnosed at a late stage after significant RGC loss and vision impairment have already occurred^5,8^, highlighting the urgent need for early detection and investigation of initial glaucomatous changes. Additionally, multiple other factors, including vascular dysfunction^9–13^, oxidative stress^14,15^, genetic predisposition^16,17^ and immune system dysregulation^18–20^, could have been implicated in complex interactions with neurodegeneration processes, complicating both mechanistic understanding and treatment development. A comprehensive understanding of glaucoma progression and its contributing biological factors could unveil new therapeutic targets and strategies for early detection and effective intervention.

Extensive research efforts have been dedicated to unravelling the pathophysiology of glaucoma. Since the retina is highly metabolically active tissue with a high demand for oxygen and nutrients, one potential cause of RGC death in glaucoma is the dysfunction of retinal blood vessels^10,11^. Though changes in retinal blood vessel morphology and ocular blood flow (OBF) are widely reported in glaucoma patients^10,11^ and animal models^9,12,21^, many previous studies utilized technologies for indirect estimation of OBF^21–24^ or mainly targeted large vessel branches such as arterioles and venules due to limited imaging resolution^11^. Studies of the finest capillaries in mouse retina under glaucoma conditions based on invasive approaches require surgery to expose the sclera, which prevents longitudinal tracing of changes in the same capillary and may cause immune system dysregulation in the retina^12,25,26^. Besides vascular dysfunction, immune system dysregulation also plays a key role in glaucoma pathogenesis. Microglia, the resident immune cells of the central nervous system, maintain retinal homeostasis under normal conditions^19,20^, but become activated in glaucoma and release pro-inflammatory cytokines, reactive oxygen species (ROS), and other neurotoxic factors, causing damage to RGCs and their axons and contributing to the progression of glaucomatous neurodegeneration^27,28^. However, results based on *ex vivo* studies cannot provide accurate information on the dynamic state changes of microglia that directly relate to their functions *in vivo*. For studying RGC activity, longitudinal functional imaging of mouse RGCs using scanning laser ophthalmoscopy (SLO) provides significant insight into the dynamic activity changes of RGCs from early to late stages of glaucoma^5^. Nevertheless, due to the large aberration of mouse eye^29^, imaging resolution for RGCs is limited to the single-cell level, making it impossible to resolve detailed structures and functions of RGCs such as axons and dendrites, which are thought to show morphologic and functional changes earlier than RGC soma degeneration^30,31^. Considering the limitations of recent glaucoma studies, a longitudinal *in vivo* high-resolution imaging technique is critical not only for early detection of glaucomatous changes but also for evaluation of treatment.

Two-photon excitation fluorescence microscopy (TPEFM) has emerged as a powerful tool offering unique advantages for *in vivo* imaging such as low phototoxicity and high imaging resolution^32,33^. With the help of adaptive optics (AO), an optical technique initially developed for astronomical telescopes, large aberrations of the mouse eye can be corrected, enabling *in vivo* imaging with subcellular resolution of mouse retina through AO-TPEFM^34,35^. Recently, a silicone oil-induced ocular hypertension/glaucoma (SOHU) model has been reported, involving the injection of a droplet of silicone oil (SO) into the anterior chamber of the mouse eye^36,37^. The SOHU model offers advantages in simplicity and faster RGC degeneration compared with other hypertension glaucoma models^8^. However, it causes additional aberration to the mouse eye, making *in vivo* high-resolution imaging more challenging^38^. In this work, we first developed an AO-TPEFM system capable of *in vivo* imaging of mouse retina under the SOHU model to achieve subcellular resolution. We then conducted a comprehensive longitudinal study of the pathological changes in retinal microvascular circulation, microglia, and RGCs during glaucomatous progression. In detail, we first captured the gradual decrease of retinal thickness during the disease through longitudinal measurements of the distance between three layers of retinal vascular plexus^39,40^. We then revealed the complex pathological alterations in flow speed and flux of retinal capillaries by tracking the blood flow dynamics of individual capillaries across three retinal vascular layers from the onset of glaucoma. With high-resolution *in vivo* imaging, we clearly captured early changes in microglial morphology and the remodeling of microglial function across different disease stages by quantifying the motion of microglial processes. Additionally, we observed the entire degeneration process of RGCs, starting from earlier dendritic degeneration followed by later soma degeneration. Furthermore, we demonstrated the importance of high-resolution functional imaging of RGCs in improving cell recognition capability and investigated the dynamic activity changes of RGCs throughout the complete degeneration process with high accuracy and fidelity.

## Results

### AO-TPEFM Enables *in vivo* Imaging in SO Injected Mouse Eye with Subcellular Resolution

We initially assessed whether ocular aberrations could be corrected in the presence of SO to achieve imaging resolution comparable to that of a normal mouse eye using our previous AO-TPEFM system^34^. After injecting SO into the anterior chamber to establish the SOHU model, we observed a distinct boundary (Fig. 1a) due to the refractive index mismatch between aqueous humor (1.336) and SO (1.405)^41,42^. When attempting AO correction in a Cx3CR1-GFP mouse using the GFP signal as a guiding star, microglial processes were surprisingly barely visible (Fig. 1b, left). To investigate this limited correction, we analyzed the focus spot patterns on our custom-built Shack-Hartmann wavefront sensor (SHWS) and found that the SO boundary’s sudden refractive index change caused excessive beam shift, hindering accurate measurement of spot positions (Fig. 1c, left). Since the laser beam covered the whole pupil of the mouse eye (larger than the typical SO diameter of ∼1.8 mm; see Methods), we optimized the system by reducing the laser spot diameter on the cornea to ∼1.3 mm (Fig. S1) and carefully aligning the beam center with the SO center. This adjustment reduced the numerical aperture (NA) from 0.458 to 0.324, preventing wavefront measurement inaccuracies and allowing clear spot images on the wavefront sensor (Fig. 1c, right). As a result, fine microglial processes became clearly resolved after AO correction with reduced NA (Fig 1b, right), though theoretical diffraction-limited resolution was reduced due to the smaller effective NA. Line profiles comparing microglial intensity between full and reduced NA correction at the same site showed resolved processes and a fourfold intensity increase with our optimized system (Fig. 1d). Further analysis of aberration amplitude in mouse eyes before and after SO injection, using peak-valley (P-V) and root mean square (RMS) values, revealed no significant difference between normal and SO-injected eyes with our reduced NA system (Fig. 1e, 1f). This suggests that significant aberration arises from the SO droplet boundary, while the aberrations induced by the droplet itself and misalignment between droplet and mouse eye centers are readily correctable.

**Fig. 1.**
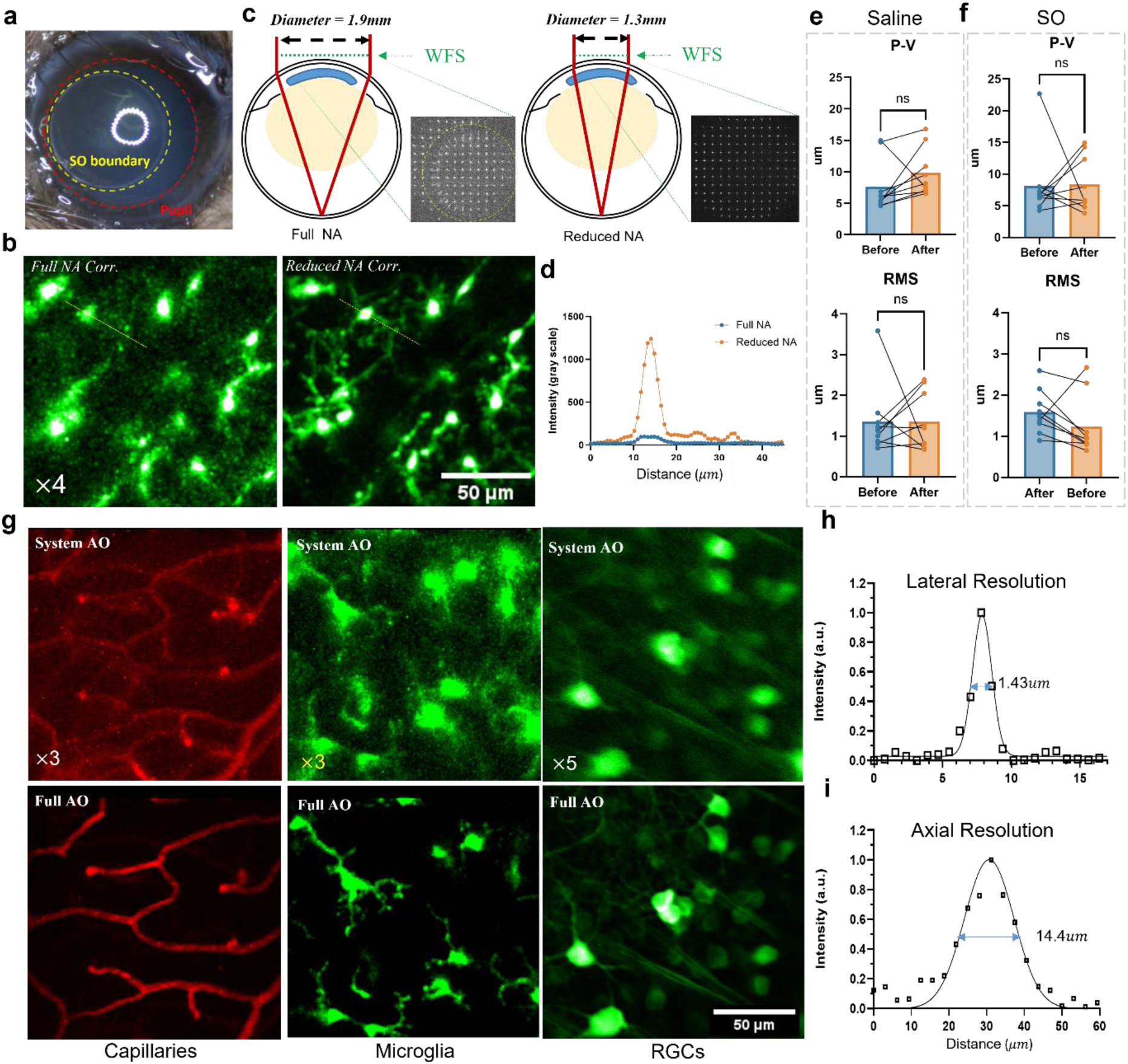
AO-TPEFM of reduced NA enables *in vivo* high-resolution imaging of SOHU mouse retina. **a.** Photo of SO injected mouse eye. The Yellow circle denotes the boundary of SO. Red circle denotes the pupil. **b.** Retinal microglia image after AO correction with full NA (left) and reduced NA AO-TPEFM (right). **c.** Schematics of error of wavefront measurement caused by the boundary of SO. Laser beam of full NA AO-TPEFM (left) has larger beam diameter than SO diameter, which causes measurement error on wavefront sensor due to the large curvature of SO boundary. Laser beam of reduced NA AO-TPEFM (right) has smaller beam diameter than SO diameter, which avoids measurement error on wavefront sensor and results in accurate wavefront measurement. **d.** Signal profiles along the dashed lines in (**c**) for a comparison of the fluorescence intensity of microglia with full NA (blue line) and reduced NA (yellow line) AO correction. **e-f.** Peak-Valley (P-V) value (top) and RMS (below) of the same eye aberration before and after saline injection (**e**) and SO injection (**f**), respectively. Aberration measured from the same eyes of 9 mice in each group. P value: 0.7344 (top), 0.6523 (below) in (**e**), P value: 0.0977 (top), 0.1289 (below) in (**f)**. Wilcoxon matched-pairs signed rank test. ns, no significance. **g.** Reduced NA TPEF images of retinal capillaries, microglia and RGCs with system (top) and full (below) AO correction. **h-i.** The lateral (**h**) and axial (**i**) resolution of the reduced NA AO-TPEFM system after AO correction. The mean values of the measured FWHM from 4 microglia are shown in the figures.

We further evaluated the reduced NA AO-TPEFM system’s performance for retinal imaging by comparing *in vivo* images of retinal blood vessels, microglia, and RGCs before and after AO correction in SO-injected eyes. Since two-photon fluorescence excitation efficiency is proportional to the fourth power of NA^43^, reducing NA decreases both excitation efficiency and imaging resolution. To counteract this efficiency loss, we optimized the pulse duration (∼100 fs) of the 920 nm femtosecond laser by compensating for system dispersion, thereby enhancing signal intensity without increasing laser power. This optimization allowed a low imaging power of just 13 mW to achieve high-quality, high-contrast images, enabling clear visualization of the finest capillaries, microglial processes, and RGC dendrites and axons after AO correction (Fig. 1g, Supplementary Video S1 and S2). To estimate *in vivo* imaging resolution, we analyzed the fluorescence intensity profiles of fine microglial processes under three conditions: full NA^34^, reduced NA without SO, and reduced NA with SO (Fig. S2 and S3)^34^. In SO-injected eyes with reduced NA, we achieved lateral and axial resolutions of approximately 1.43 μm and 14.4 μm, respectively, close to the theoretical diffraction limits (1.07 μm lateral and 14.4 μm axial calculated for 920 nm at 0.324 excitation NA). These resolution estimates are conservative, as the actual microglial process dimensions may not be much smaller than the diffraction limit. In summary, our optimized system provides sufficient imaging performance to conduct all morphological and functional imaging on SO-injected eyes^3433^ for studying glaucoma progression in the SOHU model.

### Classification of Glaucoma Progression in SOHU Model

The SOHU model was selected in this study for its straightforward operation and rapid degeneration speed^8^. The physiological condition of retina was imaged before SO injection. Following SO injection, *in vivo* imaging was performed at 0.5 weeks post SO injection (wpi), 1.5 wpi, 4.5 wpi and 7.5 wpi, adhering to a protocol as in previous longitudinal studies on SOHU model^5^. After the *in vivo* experiment at last time point, the mouse retina was extracted and stained with RPBMS to label the RGCs (see Methods). RGCs were quantified from *ex vivo* retinas in three regions (Fig. S4a): peripheral retina (over 1000 μm from optic nerve head (ONH)), mid-peripheral retina (600-1000 μm from ONH) and central retina (<600 μm from ONH). From the quantification results, we surprisingly noticed that mice were clustered into three distinct groups with significantly different RGCs survival density at three retina areas (Fig. S4a and Fig. S4b). In details, the first group of mice exhibited significant loss of RGCs in the peripheral retina, slightly loss in the mid-peripheral retina and no significant loss in the center area at 7.5wpi. We define this group as “normal-glaucoma” since the degeneration progress resembles the previous reports^5,8^. The second group showed no significant RGC loss in any of the three areas after 7.5 wpi, which is defined as “under-glaucoma” suggesting that glaucomatous progression is undetectable before this time point. The third group, labeled as “acute-glaucoma”, demonstrated rapid and severe degeneration in the peripheral and mid-peripheral areas, with significant RGC loss in the central area as early as 0.5 wpi (Fig. S4a and Fig. S4b). IOP measurements across three groups revealed no significant differences (Fig. S4c and Fig. S4d), indicating that classification cannot rely solely on IOP levels. Separating these distinct groups is crucial and reasonable due to their markedly different pathological profiles which could lead to inconsistent results if they are mixed. Unless otherwise specified, subsequent results pertain to the “normal-glaucoma” group.

### Longitudinal Imaging of Structural and Functional Changes in Retinal Microvascular Networks during Glaucoma Progression

Extensive evidence indicates that retinal thickness decreases during glaucoma progression^44–46^. This thickness, typically measured using optical coherence tomography (OCT), is considered an important biomarker for assessing glaucoma severity. For our evaluation of retinal thickness using AO-TPEFM, we leveraged the distribution characteristics of retinal vessels. Mouse retinal vasculature consists of three interconnected plexuses^39,40^: the superficial plexus (SP), intermediate plexus (IP) and deep plexus (DP) as shown in Fig. S5a. The SP is located at the surface of retina, colocalizing with the retinal ganglion cell layer (GCL). The IP is situated between the inner plexiform layer (IPL) and the inner nuclear layer (INL), while the DP is positioned between INL and outer plexiform layer (OPL)^39^. The distance between SP and DP encompasses the depth of most of the inner retina^47^ and can be used to estimate retinal thickness during the glaucoma progression, aided by the high axial resolution after AO correction. To label the retinal vessel, Texas Red (70KDa-Dextran, 100 μl, 2mg/100ul) was injected retro-orbitally into the mouse^48,49^. Retinal capillaries on different vascular layers were imaged longitudinally from before SO injection to 7.5 wpi. The same imaging site was relocated across time points by referencing large vessel branches and capillary patterns. As shown in Fig. S5b, three layers of retinal vessel were clearly visualized and the distance between DP, IP and DP could be accurately measured from the cross-sectional images of the imaging stacks (Fig. S5c). During the glaucoma progression, the distance between SP and DP shows significant decrease as early as 1.5 wpi and continued to decrease gradually, while retinal thickness remained constant in the control group with saline injection (Fig. S5c-f, Fig. S6). The total thickness decrease resulted from reductions in both the SP-IP and IP-DP distances. The more significant and earlier decrease in the SP-IP distance suggests that the disruption of connections between bipolar cells and RGCs occurs earlier in glaucoma progression^50,51^. Interestingly, in the “under-glaucoma” group, retinal thickness did not change up to 7.5 wpi (Fig. S7) despite significant IOP increases, which is consistent with the *ex vivo* results of RGC survival rates (Fig. S4a, S4b). In contrast, the “acute-glaucoma” group showed retinal thickness reduced by half as early as 0.5 wpi (Fig. S8a-e), indicating rapid and severe retinal degeneration.

OBF is a crucial and sensitive biomarker for vascular functionality. The reduction of OBF has long been considered a contributing factor to glaucomatous optic neuropathy^9–13^, but some studies report no reduction in OBF during glaucoma progression^52,53^. This inconsistency may arise from differences in disease stages, models, and the indirect methods used to estimate OBF, such as laser Doppler techniques^22,23,52,53^ and optical coherence tomography angiography (OCTA)^21,24^. Additionally, previous studies mainly focused on large retinal vascular structures, such as arterioles and venules, due to limited imaging resolution^11^. In this study, we utilized the high-resolution *in vivo* imaging capabilities of AO-TPEFM to longitudinally track the function of individual capillaries from the onset to late stages of glaucoma. This comprehensive approach aims to provide insights that may explain the discrepancies observed in earlier studies.

To study the function of microvascular circulation, we measured the blood flow speed and flux of each capillary by line-scanning along its direction^54^ (see Methods). We found that capillaries could be classified into distinct functional types. A normally functioning capillary exhibited consistent flow and flux, as indicated by the time-space image of line-scanning (Fig. 2a). However, we observed several atypical capillary functions even under physiological conditions. One type, termed “no flow,” displays a perpendicular dark band in the time-space image due to a red blood cell (RBC) becoming lodged in the capillary (Fig. 2b). Another type, “no flux,” occurred when no RBCs passed through the capillary (Fig. 2c). While these capillaries might still maintain plasma flow, we could not measure it without RBC tracers. The final type was “no perfusion/degeneration”, observed when a tracked capillary structure disappeared during a subsequent imaging time point (Fig. 2d-f). This disappearance could be due to either lack of plasma perfusion or degeneration of endothelial cells. For simplicity, the first type is referred to as “normal,” while the other three types are collectively called “abnormal” in the following sections, even though they occasionally occur under physiological conditions. The function status of individual capillaries across different vascular plexus layers were classified longitudinally before and up to 7.5 weeks post saline/SO injection. Remarkably, we found a significant increase in the percentage of abnormal functions as early as 0.5 wpi at all vascular layers (Fig. 2g). Most abnormal capillaries appearing after SO injection fell into the “no flux” and “no perfusion/degeneration” categories, indicating a severe reduction in oxygen supply to surrounding neurons. These abnormalities could result from the shrinkage of capillary diameters^12,55^. However, since diameter changes of most capillaries in glaucoma are reported to be less than 1 *μm*^12^, which is beyond our imaging resolution, we did not measure changes in capillary diameter in this work. Additionally, we observed a higher ratio of abnormal capillaries in the DP and IP compared to the SP, suggesting a possibly higher vulnerability of capillaries in the deeper plexuses in response to elevated IOP. We also analyzed the changes in flow speed and flux of capillaries across the three vascular layers in the under-glaucoma group. We found no difference in the distribution of normal and abnormal capillaries between the control and experimental groups (Fig. S9c), indicating that most capillaries functioned normally after SO injection. In contrast, the acute-glaucoma group showed a severe loss of capillaries at 0.5 wpi (Fig. S10c), demonstrating strong degeneration of retinal vascular structure and function in this condition.

**Fig. 2.**
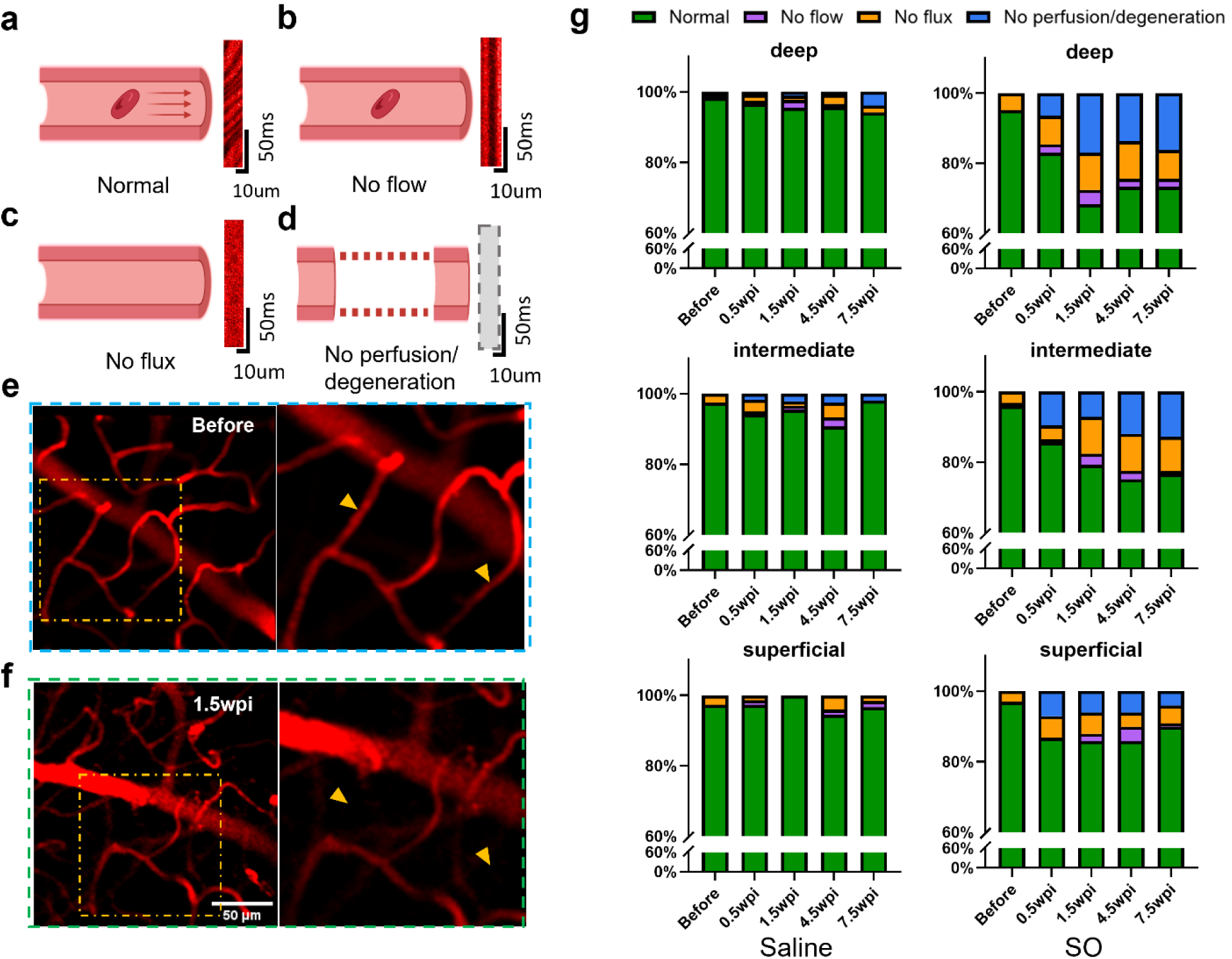
Functional changes of retinal capillaries during the progression of glaucoma. **a.** Schematics of capillary that has normal function. Both flow and flux exist. Right figure shows the image of line-scan along the capillary. **b.** Schematics of capillary that has no flow speed. Right figure shows the line-scan image of a RBCs get stuck in the middle of a capillary. **c.** Schematics of capillary that has no flux. Right figure shows the line-scan image of a capillary without any RBCs passing by. **d.** Schematics of capillary that disappear. There are two possibility for this observation: no perfusion or degeneration of capillary endothelium cell. **e**-**f.** Illustration of schematics in (**d)**. Longitudinal tracing of same capillaries shows that two capillaries (yellow arrow head) exist before SO injection (**e**) but disappear at 1.5 wpi (**f**). **g.** Contingency table of the ratio of 4 types of capillaries defined in Fig **a-d** at different time point after saline/SO injection (6 mice in control group, 6 mice in experiment group). Left column denotes the ratio of 4 types of capillaries at different vessel layers after saline injection. Right column denotes the ratio of 4 types of capillaries at different vessel layers after SO injection. Figure (**a-d**) created using BioRender (https://biorender.com/)

Next, we conducted a quantitative study of blood flow speed and flux of capillaries during glaucoma progression. We first grouped all capillaries into 2 sub-types based on their functional behavior across all imaging sessions. Capillaries that consistently exhibited normal function throughout all imaging sessions were classified as sub-type 1 (Fig. 3a), suggesting they were less likely to be affected by glaucoma. Capillaries that displayed any of the other three abnormal functions at least once during any imaging session were classified as sub-type 2 (Fig. 3b). After dividing capillaries into these two groups, we compared the flow speed and flux between sub-types 1 and 2. Since the flow speed is zero for “no flow” capillaries and the flux is zero for “no flux” capillaries, we excluded these zero values from the flow speed and flux data of sub-type 2 in the subsequent analysis. This exclusion prevented the mean values of flow speed and flux from being artificially skewed toward lower values.

**Fig. 3.**
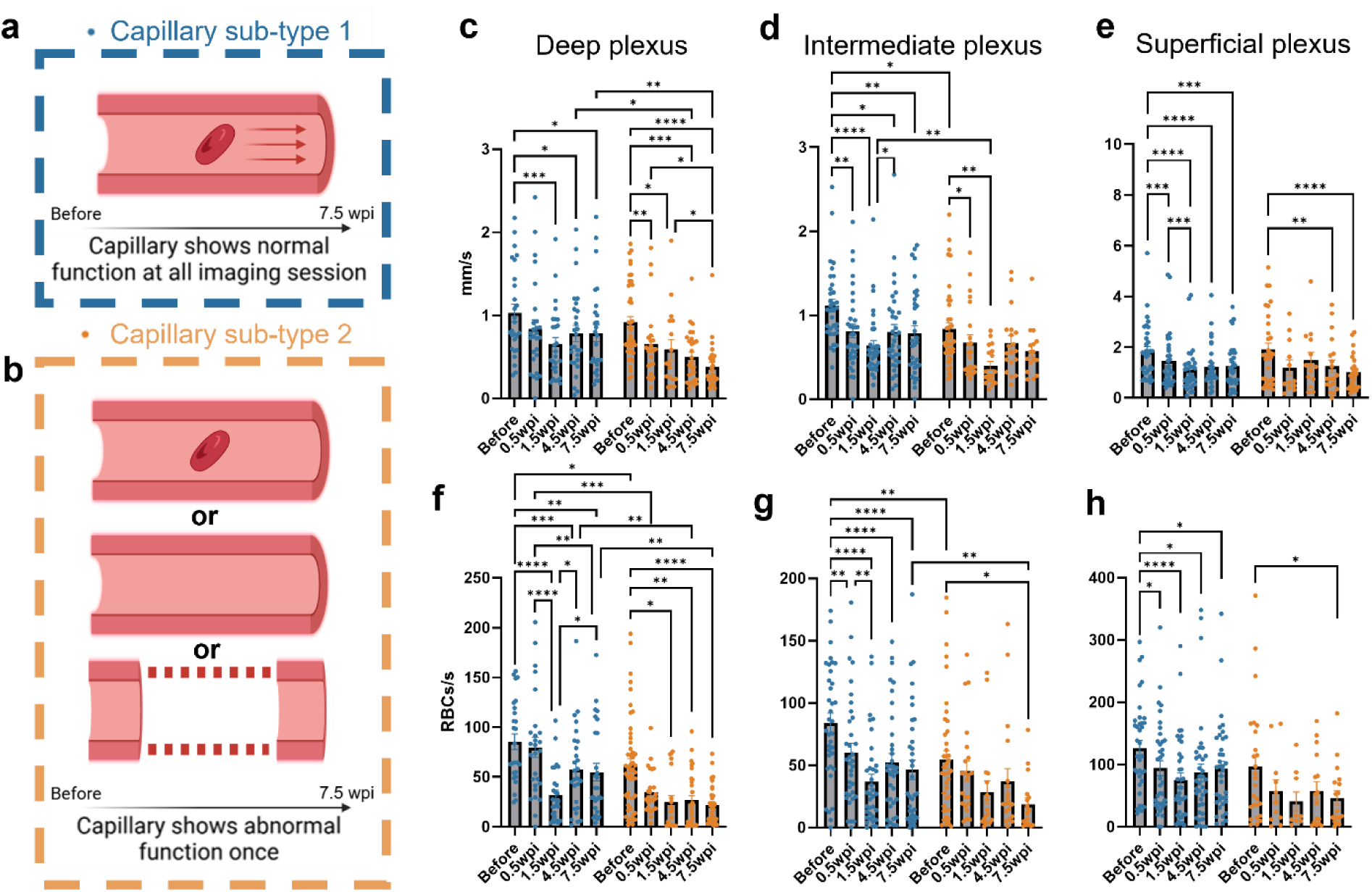
Flow speed and flux of sub-type 1 and sub-type 2 capillaries during progression of glaucoma. **a**. Illustration of capillary sub-type 1, which is defined as the capillary that shows normal function at all imaging sessions. **b.** Illustration of capillary sub-type 2, which is defined as the capillary that shows no flow, no flux, or no perfusion/degeneration once during all imaging sessions. **c-e.** Flow speed of individual capillary at different retinal vessel layers of two sub-types at different time point after SO injection (sub-type 1: blue; sub-type 2: yellow). **f-h.** Flux of individual capillary at different retinal vessel layers of two sub-types at different time point after SO injection (sub-type 1: blue; sub-type 2: yellow). Data in (c-h) were present as mean value with individual data point. Mixed-effects model analysis with Tukey’s multiple comparisons test, ns is not shown. *p<0.05; **p<0.01; ***p<0.001; ****p<0.0001. Data obtained from 6 mice of normal-glaucoma group. Figure (**a-b**) created using BioRender (https://biorender.com/)

In general, there are similar patterns between blood flow speed and flux, and the initial flow speed and flux of sub-type 2 was slightly lower than that of sub-type 1 (Fig. 3c-h). In addition, a reduction in blood flow speed and flux following SO injection is observed in both sub-type 1 and sub-type 2 capillaries, whereas the flow speed and flux remain quite stable following saline injection over the 7.5-week period (Fig. S9a and S9b). Specifically, in the DP, sub-type 1 capillaries show a significant decrease at 1.5 wpi, with partial recovery observed at 4.5 wpi and 7.5 wpi. Surprisingly, sub-type 2 capillaries experience a continuous, monotonic decrease in blood flow speed starting from 0.5 wpi to 7.5 wpi (Fig. 3c). Similar patterns are also observed in the flux at DP (Fig. 3f). Additionally, the significant reduction in flow speed and flux for sub-type 2 occurs earlier (0.5 wpi) compared to sub-type 1 (1.5 wpi).

In the IP and SP, both sub-type 1 and sub-type 2 exhibit obvious reductions earlier at 0.5 wpi, followed by a slight recovery at 4.5 wpi or 7.5 wpi (Fig. 3d-e and g-h). No obvious difference of change patterns of flow speed and flux was observed between sub-type1 and sub-type 2 in IP and SP compared with DP. The average flow speed and flux were significantly higher in the SP compared to the DP and IP (Fig. 3e and h). This is because large arterioles were located in the SP and branched into the DP and IP, making it more likely to measure capillaries closer to the upstream arterioles^39^. Considering the more obvious increase in the ratio of abnormal function capillaries in DP (Fig. 2g) and a monotonic decrease of blood flow speed and flux of capillaries in DP sub-type 2, DP capillaries might be more vulnerable compared with IP and SP capillaries. A possible explanation is that DP capillaries are furthest away from the large arterioles in SP which results in a lower blood pressure and causes a more susceptible hemodynamics in response to increased IOP.

In the under-glaucoma group, we analyzed flow patterns differently. Due to the limited number of capillaries exhibiting abnormal function in this group, we compared the flow speed and flux of all capillaries collectively, rather than separating them into sub-type 1 and sub-type 2 (Fig. S9c). Contrary to observations in the normal-glaucoma group, we noted a slight increase in flow speed at the early stages (0.5 wpi) across all three vascular layers following SO injection, although this gradually decreased by 7.5 wpi (Fig. S9a and S9b). The flux exhibited a much slower decrease in the DP and IP compared with the normal-glaucoma group, showing significant changes only at 7.5 wpi, while remaining quite stable in the SP. The initial increase in blood flow and flux may play a significant role in the observed resistance to glaucomatous changes in the under-glaucoma group. In contrast, the acute-glaucoma group showed a substantial reduction in blood flow speed and flux following SO injection, indicating marked degeneration of retinal vascular function and distribution (Fig. S10).

### Dynamic Morphological and Functional Changes of Microglia during Glaucoma Progression

Microglia, the primary immune cells of the CNS, play a critical role in the neurodegeneration and inflammation associated with glaucoma^19,20^. Previous *ex vivo* studies have demonstrated the morphological changes of microglia^56–58^ within the glaucomatous retina, while complementary *in vivo* investigations have documented gradual alterations in microglial density^59^. Because microglia project highly motile processes that rapidly respond to damage or disease^60,61^, longitudinal *in vivo* imaging at subcellular resolution is crucial for elucidating their functional role throughout glaucoma progression. In this study, we injected SO into the *Cx3Cr1-GFP* mouse eye to image the dynamic changes of microglia throughout the glaucoma progression (Fig. 4). Retinal vessels were labeled as landmarks to ensure consistent relocation of the same imaging sites during each session (Fig. 4a).

**Fig. 4.**
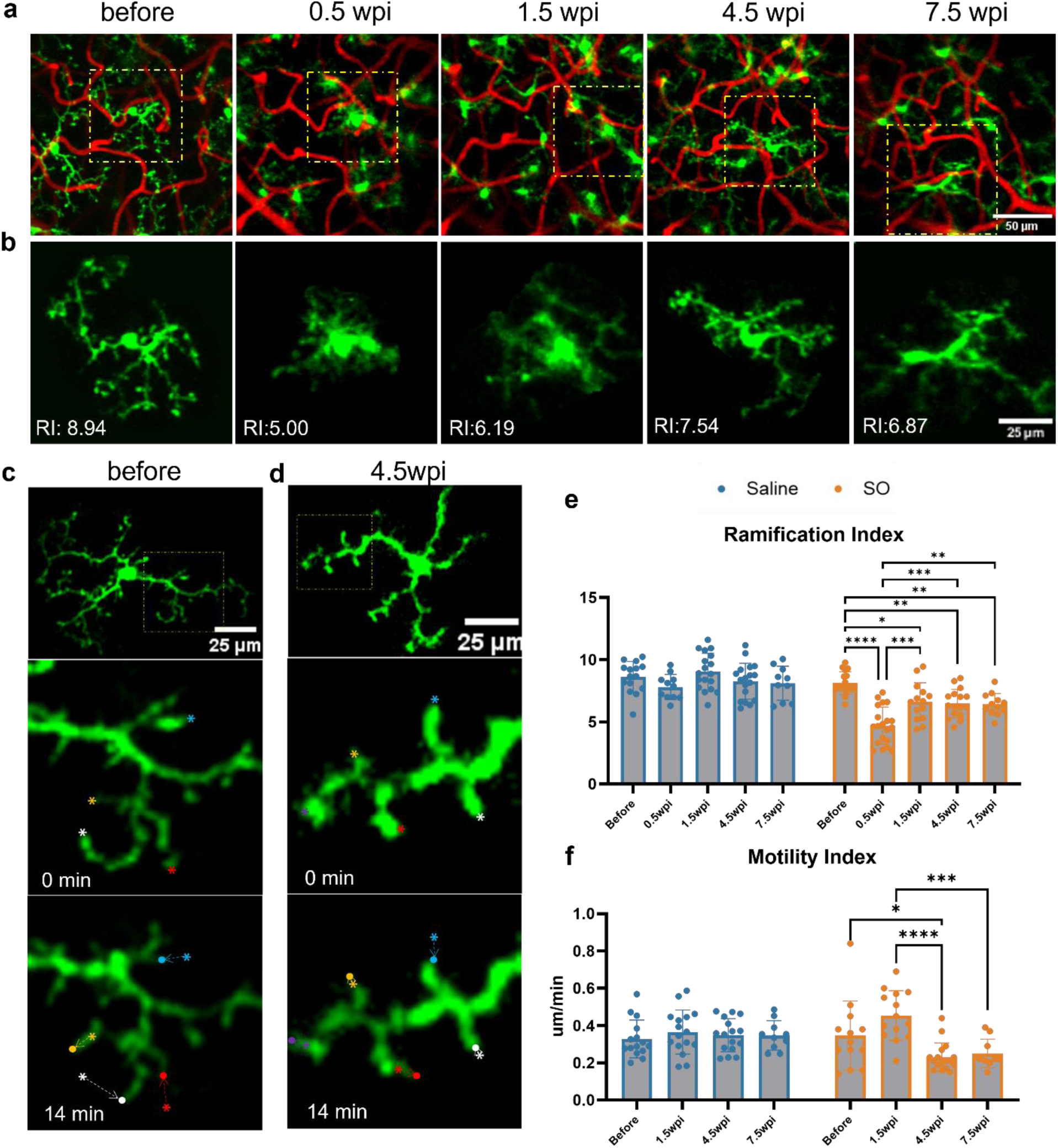
Morphological and functional changes of retinal microglia during the progression of glaucoma. **a.** Longitudinal recordings of blood vessel structures (red) and microglia (green) on a same site in retina at different time point. **b.** Small FOV image of the microglia in the square area with yellow dashed line in **a**. Ramification index of each microglia is shown on the left corner for each microglia. **c.** Time-lapse images of the dynamic processes of microglia before SO injection. Below images are the small FOV of the microglia processes movement in the square are with yellow dashed line. Initial position of different process tips is labelled with asterisk with different colors and the final position of the process after 14mins is labelled with dot of corresponding color. **d.** Time-lapse images of the dynamic processes from microglia after 4.5 weeks post SO injection from the same mouse. Below images are the small FOV of the microglia processes movement in the square are with yellow dashed line. Initial positions of different process tips are labelled with asterisk with different colors and the final position of the process after 14 mins is labelled with a dot of corresponding color. **e-f.** Statistics of microglia ramification index (e) and motility index (f) at different time points. 3 mice in control group, 2 mice in glaucoma group. Data was present as mean value with SD and individual data point. Two-way ANOVA with Tukey’s multiple comparisons test. ns and comparison between control and experiment group are not shown. *p<0.05; **p<0.01; ***p<0.001; ****p<0.0001.

In this study, significant microglial activation was detected as early as 0.5 wpi, characterized by a reduction in the ramification index (RI)^62^, a quantitative measure of microglial morphology (Fig. 4b). The control group maintained stable RI values throughout the observation period (Fig. 4e and Fig. S11), indicating that microglia retained their normal surveillance morphology following saline injection. Interestingly, from 1.5 weeks post-SO injection, the RI showed a slight recovery, although it remained lower than in the control group (Fig. 4e). To complement our morphological analysis, we quantified microglial process dynamics by measuring the movement speed of process tips^60,63^, defined as the motility index (MI) (see Methods). The protocol involved acquiring sequential imaging stacks of individual microglia over 20-30 minutes intervals, followed by manual tracking of process tip positions across frames (Fig. 4c-d). We found that at 0.5 wpi, microglia exhibited high-level activation with predominantly amoeboid morphology and few remaining processes, rendering MI measurements unreliable at this timepoint and necessitating their exclusion from analysis. Longitudinal MI assessment revealed stable values in control animals throughout the study (Fig. 4f). In contrast, the experimental group demonstrated biphasic changes in MI during glaucoma progression: a transient increase at 1.5 wpi followed by significant decreases at both 4.5 and 7.5 wpi (Fig. 4d and f). These temporal alterations in process dynamics suggest functional remodeling of microglia across different disease stages, despite the relatively consistent microglial morphology observed between 1.5-7.5 wpi. This dissociation between morphological appearance and dynamic behavior highlights the importance of functional assessment in understanding microglial responses to glaucomatous damage. Future investigations could employ spatial single-cell RNA sequencing technology to elucidate the transcriptional profiles underlying the observed temporal changes in microglial dynamics^64,65^, potentially revealing stage-specific molecular signatures that drive functional transitions during glaucomatous progression.

It should be emphasized that physiologically microglia are found predominantly in the synaptic layers OPL and IPL^66,67^. Our high-resolution *in vivo* imaging also revealed layer-specific differences in microglial morphology and dynamics (Fig. S13c-d, Supplementary Video S1). However, progressive retinal thinning (Fig. 2c), compromised our ability to reliably distinguish microglia by layer at advanced disease stages. Consequently, we analyzed all microglia collectively without layer-specific comparisons of RI and MI. In addition, we found that in the under-glaucoma group, microglia exhibited transient activation at 1.5 wpi with reduced RI (Fig. S12, Fig. S13a), but recovered to normal at later timepoints. Notably, MI remained stable throughout the observation period in this group (Fig. S13b). The acute-glaucoma group demonstrated pronounced microglial activation at 0.5 wpi (Fig. S8a, S8f), with such extensive process retraction that MI analysis was not feasible at this timepoint.

### Subcellular Longitudinal Imaging Captures Accurate Dynamic RGC Degeneration in Glaucomatous Progression

To investigate RGC morphology and function, we used a Brn3b-Cre transgenic line with RGC-specific expression^68^ (Fig. S14), and labeled these RGCs with the high-sensitivity genetically encoded calcium indicator GCaMP8s^69^ via intravitreal AAV2-loxp-GCaMP8s injection. Our *in vivo* imaging focused on the ventral retina due to its balanced expression of short-wavelength (360 nm) S-opsin and middle-wavelength (508 nm) M-opsin compared to the dorsal region^70^. For visual stimulation, we employed a 405 nm UV light source^71^ to activate mouse cones, approximately 95% of which co-express both S-opsin and M-opsin^5,70,72^. ^71^To prevent UV light interference during GCaMP signal detection^73^, we synchronized stimulation to occur only during microscope scanner flyback periods. TPEFM offered significant advantages over scanning laser ophthalmoscopy (SLO) for RGC functional studies^71^, as its 920 nm excitation wavelength falls outside the mouse cone sensitivity range, minimizing visual crosstalk^34,74^. We observed that initiating the 920 nm laser triggered transient responses in some RGCs (Fig. S15a), likely due to nonlinear absorption^74^, so we implemented a 10-second delay between laser scanning initiation and stimulus presentation since these responses typically subsided within 3 seconds (Fig. S15a).^74^ The stimulation light was maintained at constant intensity (1 μW) for 20 seconds to generate robust Ca^2+^ signals that captured GCaMP8s dynamics^5,69^, with each calcium imaging session lasting 60 seconds per retinal site. For analysis, we calculated the time-dependent fluorescence change (*ΔF*) by subtracting the mean baseline fluorescence during the 10-second pre-stimulus window (*F₀*) from the measured signal (*F*), and the normalized responses (*ΔF/F₀*) were then processed through a low-pass filter for subsequent analysis.

We first evaluated the necessity of high-resolution imaging in RGCs function study. To analyze GCaMP8s fluorescent signals in naïve mouse retina, we recorded time-lapse movies and extracted single-cell signals by manually delineating RGC soma boundaries from averaged-intensity projections. Sub-cellular resolution imaging allowed for clear visualization of the boundaries between RGC somas and dendrites, enabling accurate identification of closely adjacent RGCs without merging them when the aberration is corrected (Fig. S16a and S16c, Supplementary Video S2). This precision is crucial for studying RGC functions and classifying RGC types, as low-resolution imaging can blur RGC boundaries, potentially mixing distinct types of RGC Ca^2+^ responses (Fig. S16b and S16d). Moreover, the enhanced imaging contrast with AO correction improves the dynamic range of detectable RGCs as many RGCs with low GCaMP8s signal levels might otherwise be overlooked without AO correction (Fig. S17). Having evaluated the advantages of AO-TPEFM in accurately studying RGC morphology and function, we proceeded to investigate RGC changes in response to glaucoma. Significant morphological changes in RGCs were observed following SO injection compared to saline-injected mice (Fig. 5a, Supplementary Video S3 and S4). Through longitudinal recording of the same RGCs, we discovered that RGC dendrites are lost earlier than their somas (Fig. 5a, Fig. S18), similar to previous work with early loss of axons and shrinkage of dendrites in an optic nerve crush model^75^, suggesting the possibility of initial malfunction at the RGC input during glaucoma progression.

**Fig. 5.**
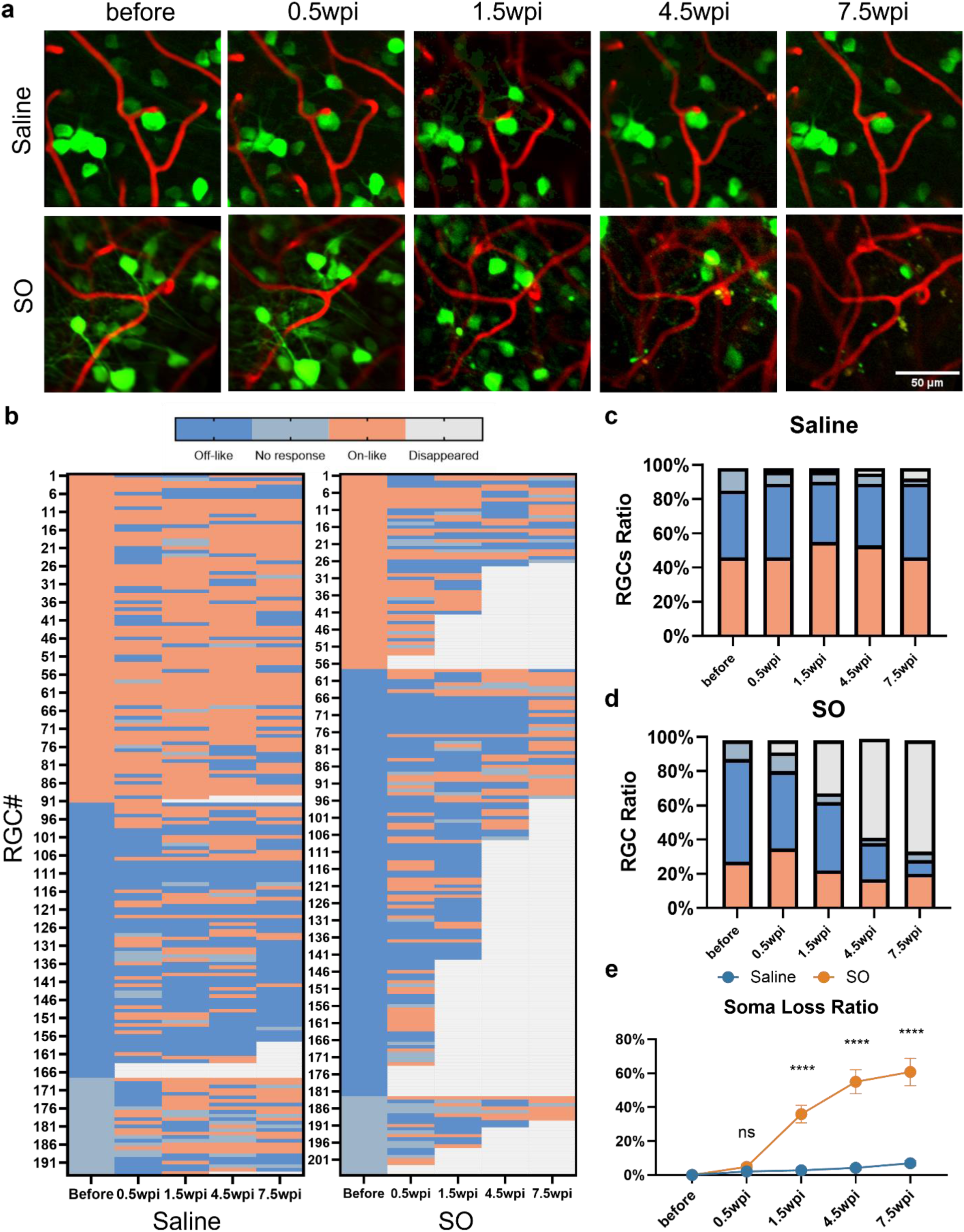
Morphological and functional changes of retinal ganglion cells during the progression of glaucoma. **a.** Longitudinal recordings of blood vessel structures (red) and RGCs (green) on a same site in retina at different time point with saline injected eye (above row) and with SO injected eye (below row). Images shown as average intensity projection of 240 frames in 60s. **b.** Heatmap of the type of each RGC recorded longitudinally at different time point after saline/SO injection. Each row represents one RGC. 4 mice for each group. The left panel denotes types of each RGC after saline injection. Right panel denotes types of each RGC after SO injection. **c.** Contingency table of ratio of different types of RGCs at different time points after saline injection. **d.** Contingency table of ratio of different types of RGCs at different time point after SO injection. **e.** Ratio of RGC loss at different time points after saline/SO injection. Data were present as mean ± SEM. Two-way ANOVA with Tukey’s multiple comparisons test. ****p<0.0001. 11 sites from 4 mice in control group, 11 sites from 4 mice in experiment group.

In glaucoma, numerous studies have observed that RGCs exhibit type-specific vulnerability^30,76,77^, meaning certain types of RGCs degenerate faster and earlier than others. These studies mainly focus on *ex vivo* results without revealing the dynamics of RGC function. Recently, a longitudinal *in vivo* study of RGC function has revealed activity changes during glaucoma progression^5^, indicating the importance of *in vivo* imaging for detecting dynamic changes in RGCs. Here, we studied RGC soma function changes with AO-TPEFM *in vivo*. While it is technically feasible to study dendrite function as well, the complex distribution of the dendrite network makes it challenging to assign each dendrite to its corresponding soma without sparse labeling of RGCs^71^. Therefore, this study focuses solely on soma function. We initially determined the range of non-specific Ca^2+^ responses resulting from spontaneous activity or imaging procedures, unrelated to UV stimulation, as described in previous work^5^. We recorded 60 s videos of RGC Ca^2+^ responses without UV stimulation and divided the RGCs into 2 groups based on whether their fluorescence signal increased or decreased between 10 and 30 seconds (see Methods). The upper and lower bounds of the non-specific range were calculated by averaging the waveforms of each group and selecting the maximum absolute value of the averaged waveform (Fig. S15b). The range of non-specific responses was found to be between −19.1% and 26.2%. We define an RGC as having no response (NR) if its maximum or minimum response value fell within this range during UV stimulation. For RGCs with responses outside this range, we classified them into two simple types: “on-like” and “off-like”. This classification was based on whether the mean value of *ΔF/F* during the stimulation period (10 s to 30 s) was greater or less than the mean value of *ΔF/F* from 5s to 10s (see Methods). The average waveforms of “on-like” and “off-like” RGCs are shown in Fig. S15c, and the combined percentages of “on-like,” “off-like,” and “no response” RGCs are presented in Fig. S15d.

Having defined three types of simple clusters of RGCs, we conducted a longitudinal study of the functional changes in RGCs during the glaucoma progression. Interestingly, even in the control group, some RGCs switched between “on-like” and “off-like” types (Fig. 5b). However, the overall ratio of “on-like” to “off-like” RGCs, as well as the total number of RGCs, remained stable up to 7.5 wpi (Fig. 5c). Additionally, some “no-response” RGCs exhibited calcium responses at different time points, although the overall proportion of “no-response” RGCs remained generally unchanged. In the experimental group, the functional changes in RGCs were markedly different, with RGC disappearance beginning at 0.5 wpi and progressively worsening at later stages (Fig. 5b and 5d-e). Notably, there was a significant functional conversion in RGC activity following SO injection, particularly among “off-like” RGCs. Although there was a significant loss of “on-like” RGCs during the disease progression, most surviving “off-like” RGCs converted to “on-like” RGCs by 7.5 wpi, resulting in a stable survival rate for “on-like” RGCs (Fig. 5b and 5d). Future studies may leverage spatial single-cell RNA sequencing to characterize transcriptional landscapes associated with the functional conversion in RGCs^78^, potentially identifying specific molecular mechanisms that orchestrate functional transitions related to survival capability of RGCs during glaucoma progression. Similar to the control group, no obvious morphological changes were observed up to 7.5 wpi, in the under-glaucoma group (Fig. S19a). However, there was a gradual increase in the ratio of “on-like” RGCs starting from 0.5 wpi (Fig. S19b-d). In contrast, the acute-glaucoma group exhibited significant morphological changes in RGCs, with numerous debris and broken capillaries observed as early as 0.5 wpi (Fig. S20), consistent with previous results of severe loss of vascular function and strong activation of microglia.

## Discussion

As a complex neurodegeneration disease with interplay among various factors, glaucoma has been studied for decades using different approaches such as *ex vivo*^7,56,57^, *in vivo*^5,12,59^ or gene expression analysis^17,79^, yet the mechanism of glaucoma has remained elusive. An *in vivo* longitudinal study with high resolution from early to late stages of glaucoma would provide a good opportunity for understanding the disease progression and evaluating treatment effects. In this study, we demonstrated the feasibility of longitudinal *in vivo* imaging of the mouse retina during glaucoma progression at sub-cellular resolution using optimized AO-TPEFM on the SOHU model. We found that the boundary of the SO introduces prominent aberration, leading to incorrect focal shift measurements in the WFS (Fig. 1b). This can be easily understood through two lenses with different focal lengths and diameters, placed coaxially and adhesively. When the laser beam diameter is smaller than both lens diameters, the lenses function as a simple doublet with a fixed focal length. However, if the beam diameter exceeds the smaller lens, light beams beyond its boundary result in a different focal length, causing a sudden change in beam direction at the smaller lens boundary. In our current AO-TPEFM setup, this distortion is too large to be accurately measured by the WFS. To address this, we reduced the beam size to avoid measurement errors. Optimizing the microlens array to increase the detection angle of the input wavefront is another potential solution for managing large distortion angles without sacrificing numerical aperture (NA), an approach we could explore in future work.

As reduced NA decreases the two-photon excitation efficiency ^43^, we optimized the laser pulse duration to approximately 100 fs by compensating the system dispersion to enhance two-photon excitation efficiency instead of increasing laser power which has a high risk of causing laser damage^80,81^. Due to the varying pixel exposure times between line scanning of capillary (<20 μm, 1 KHz line rate), small FOV guiding star measurement (<10 μm × 10 μm, 2 Hz frame rate), and large FOV imaging (<200 μm × 200 μm, 2-4 Hz frame rate), the safety thresholds for laser power differ accordingly^82^. We empirically applied different imaging powers for each scanning mode (see Methods) and assessed the safety of these power parameters by longitudinally monitoring microglial morphology and dynamics in control groups at various time points (Fig. 3e and 3f). Our observations revealed negligible changes in RI and MI across multiple imaging sessions in control groups, suggesting that the imaging power is safe for longitudinal *in vivo* imaging on the pigmented (C57BL/6J background) mice^83–86^. We did not investigate whether higher safe imaging power could be used in albino mice, which have been reported to be more resistant to laser damage due to reduced absorption by the retinal pigment epithelium (RPE)^80^. However, given the sufficient signal intensity achieved with our current dye concentration (Texas Red) and fluorescent protein expression levels (GFP and GCaMP8s), albino mice could be a viable option for future studies requiring higher imaging power for fluorescent proteins or dyes with weaker signal intensity.

Among the various models of glaucoma, the SOHU model has its advantages in simplicity and rapid degeneration speed^8,36^. In our study, we identified three distinct groups based on the rate of degeneration following SO injection, which we termed “normal-glaucoma,” “under-glaucoma,” and “acute-glaucoma”. It is important to note that our classification differs from previous study^36^ where the degeneration speed may be modulated by the pupil dilation frequency. In this work, pupil dilation was performed only at 0.5 wpi, 1.5 wpi, 4.5 wpi, and 7.5 wpi before *in vivo* imaging, similar to the protocol of previous work^5^. Specifically, the “normal-glaucoma” group exhibited the most similar result to previous reports ^5,8,36^ regarding the degeneration of RGCs at 7.5 wpi. In contrast, the “under-glaucoma” group showed rather slow and minimal changes in the morphology and function of retinal vessels, microglia, and RGCs. The “acute-glaucoma” group experienced severe degeneration, as indicated by all biomarkers in less than 1 week, resembling acute glaucoma after several months following SO injection in patients^87^. Interestingly, IOP did not significantly differ among these groups (Fig. S4c and S4d), suggesting a complex underlying mechanism. One possible explanation is the stability of high IOP after SO injection, as it has been proved that a high and stable IOP can increase the RGC degeneration speed^36^. Although the IOP was similar between each group at each imaging time point, fluctuations in IOP between imaging sessions could have been missed, potentially influencing degeneration speed.

A notable discrepancy from the original SOHU model is the distribution of the three identified groups. From all our *in vivo* imaging mice with SO injection (n = 38), only a small proportion (n = 8, 21.0%) exhibited characteristics of the “normal-glaucoma” group. In contrast, the majority of mice (79.0%) fell into the other two unexpected groups, with 60.5% (n = 23) classified as “under-glaucoma” and 18.5% (n = 7) as “acute-glaucoma.” Despite meticulously replicating the SO injection procedures, the lower percentage of mice in the “normal-glaucoma” group may be attributed to differences in imaging protocols between our AO-TPEFM and previous studies using SLO and OCT^5,8^. Our imaging protocol (see Methods) requires precise alignment of the laser beam with the SO and subsequent relocation of three imaging sites, which are time-intensive procedures. During imaging, any positional shift of the SO necessitates realignment of the laser, repositioning, and aberration correction, further extending the experiment duration. Typically, over two hours are needed to complete *in vivo* imaging after full pupil dilation. It remains unclear whether this prolonged imaging session inhibits the accumulation of IOP, as it has been established that IOP is released in the SOHU model when pupils are dilated for imaging^87^.

Morphological and functional changes in retinal vasculature have been extensively documented in glaucoma on patients^10,11^ and animal models^9,12,21^. However, most previous studies estimate the blood flow speed of large retinal vessels using indirect measurements such as laser Doppler technologies or OCTA. A recent work utilized two-photon microscopy to directly measure RBC flux changes by tracking single RBC movements within a capillary during glaucoma^12^. Although this approach provides higher resolution with a high NA objective, it is partially invasive, requiring the surgery of sclera^12,25^, and is unsuitable for long-term longitudinal tracking of the same capillary. Additionally, imaging from the sclera requires unpigmented mice (albino)^12,25^ to prevent strong absorption by the RPE. Furthermore, the low frame rate (40 Hz)^12^ 2D recording of a capillary flux limits the maximum measurable flux and flow speed given the significant variability in retinal capillary flow^88^. In this work, we achieved non-invasive direct measurement of capillary flux and flow speed using high-speed line scanning (1-2 KHz), which is fast enough to accommodate the diverse velocity levels of nearly all capillaries^88^. This technique allows for tracing and measuring each capillary across different vessel plexus layers from the onset to the late stages of glaucoma and is applicable to all types of mice. Consistent with most previous reports, both flow speed and flux generally decrease after glaucoma onset. However, when categorizing capillaries into two sub-types based on observed abnormalities, we discovered differing patterns of blood flow and flux reduction at various plexus layers during glaucoma progression. Specifically, sub-type 1 capillaries in DP and IP shows a recovery of flow speed at the late stage of glaucoma while the sub-type 2 capillaries show a more monotonic decrease, suggesting a possible different remodeling mechanisms such as the density of pericytes on each capillary^12,55,89,90^. Moreover, since the SP capillaries exhibit less severity compared to the DP and IP, it is worth evaluating whether capillaries with higher initial blood flow and flux are more resistant to the effects of OBF in future studies.

As the primary immune cells in the CNS, microglia play crucial roles in the neurodegeneration process of glaucoma^19,20^. With the help of sub-cellular resolution imaging, we observed for the first time the dynamic changes of microglial morphology and process motility at various stages of glaucoma. Interestingly, we observed initial microglial morphological alteration as early as 0.5 wpi followed by a slight recovery at later stages, indicating a remodeling of retina conditions shortly after SO injection. Notably, significant morphological alterations of microglia were observed even in the “under-glaucoma” group at 1.5 wpi (Fig. S13a), when changes in vessel function and RGCs were not detected. This finding implies that microglia may serve as the most sensitive biomarkers for indicating glaucoma, compared to vessel and RGC changes. Moreover, using *in vivo* functional imaging with high-resolution, we revealed that microglia exhibited distinct dynamic motility across various stages of glaucoma, suggesting that their functions may vary depending on the stage of the disease. Given that microglia could be either protective or destructive at different disease stages^91^, the observed transitions in microglial morphology and function suggest the possibility that microglia also show dual effects in glaucoma^92^ which warrants further evaluated in the future. Some reports show that the ischemia or oxygen-glucose deprivation may affect the motility of microglia in brain^93,94^. Considering the observed decrease in flow speed and flux in the SO group, a plausible explanation for the reduced MI at 4.5 and 7.5 wpi is the chronic reduction in oxygen supply to the glaucomatous retina at later stages.

It is now understood that RGCs are not simple light detectors but rather feature detectors that comprise various feature channels^95^. Different types of RGCs can be sensitive to different patterns such as orientation, moving direction, contrast, brightness and color^96,97^. Based on Ca^2+^ responses to light stimulation, RGCs have been classified into over 30 types from an *ex vivo* study^71^ and 9 types *in vivo*^5^. These studies are based on large sample sizes (over 8,000 RGCs), providing a reliable approximation of the actual probability distribution. In this work, due to the limited number of RGCs (fewer than 1,400), we classified only two basic types to minimize the risk of misclassification. Enhancing the throughput for high-resolution RGC recording and incorporating sophisticated stimulation pattern^71^ would be crucial for achieving accurate and reliable *in vivo* classification of RGCs in the future.

Various hypotheses have been proposed to explain the potential mechanisms underlying the type-specific vulnerability of RGCs in glaucoma, including differences in receptor expression, vascular perfusion, and RGC metabolism^76^. In this study, we also observed a higher vulnerability of off-like RGCs during glaucoma progression, with a significant likelihood of functional transition^5^ and increased degeneration ratio^30,31^. Moreover, we noticed a higher initial percentage of off-like RGCs in the normal-glaucoma group prior to SO injection (Fig. 5b), while the percentage of under-glaucoma is quite similar to control group. It should be emphasized that all mice were selected randomly, without bias. One possible explanation is that mice with a greater proportion of off-like RGCs may be more susceptible to glaucoma development. However, due to the limited number of recorded RGCs and the small proportion of mice in the normal-glaucoma group (21.0%), this hypothesis requires further investigation with a glaucoma model yielding higher success rates, and an enhanced throughput high-resolution RGC recording technology.

## Methods

### Animals

All animal procedures in this study were conducted in accordance with the guidelines of the Animal Care Facility at the Hong Kong University of Science and Technology (HKUST) and were approved by the Animal Ethics Committee of HKUST. All the animal experiments were conducted on adult male mice aged 2 to 4 months, weighing 20 to 35 grams, from the following strains: 1) C57BL/6J wild-type mice, utilized in experiments involving capillary visualization via retro-orbital injection with Texas Red; 2) green fluorescent protein-expressing mice under the control of the Cx3cr1 promoter (Cx3Cr1^GFP/+^), which enable selective visualization of retinal microglia (B6.129P2(Cg)-Cx3cr1tm1Litt/J; The Jackson Laboratory); and 3) Brn3b-iCre mice, generated using a standard CRISPR-Cas9 approach on the C57BL/6J background, with the CreER cassette inserted downstream of the first exon of the Brn3b open reading frame. Founders underwent PCR screening, and the genotypes of the F1 offspring were verified using both PCR and sequencing. This strain was subsequently backcrossed to C57BL/6J. The animals were habituated in a controlled environment with a 12:12-hour light-dark cycle, and provided with ad libitum access to food and water.

### Constructs and AAV production

AAV serotype 2/2 was used for the expression of jGCaMP8s (Addgene plasmid # 162380). The AAV titers were determined by real-time PCR and diluted to 5×10^12^ vector genome (vg)/ml for intravitreal injection.

### Intravitreal injection and tamoxifen injection

The following reagents were administered by intravitreal injection (2 μl total volume): pAAV2/9-CAG-FLEX-jGCaMP8s-WPRE or PBS. The tip of a custom-made glass micropipette was inserted into the superior quadrant of the eye at an approximately 45° angle, through the sclera into the vitreous body, avoiding injury to eye structures or retinal detachment. Antibiotic ointment was applied upon the cornea to prevent bacterial infection and promote wound healing. 14 to 30 days after virus injection, tamoxifen (75 mg/kg once per day for 5 days) was intraperitoneally injected. Specific expression of GCaMP8s after tamoxifen treatment was evidenced by co-localization of the pan-RGC marker RBPMS and GCaMP8s in the inner retina layers (Fig. S14a and S14b) and no amacrine cell is labelled with GCaMP8s (Fig. S14c). The mice were housed for an additional 2 weeks to achieve stable jGCaMP8s expression.

### Immunohistochemistry and RGCs counts

Mice were anesthetized and subjected to transcardial perfusion with 4% paraformaldehyde (PFA). The eyes were then removed and post-fixed in 4% PFA for two hours at room temperature. For the preparation of retinal wholemounts, the retinas were dissected, thoroughly washed in PBS, and blocked in a staining buffer containing 10% BSA and 2% Triton X-100 in PBS for 30 minutes. Primary antibodies used included Rabbit anti-RBPMS (Phosphosolutions), Chicken anti-GFP (Invitrogen) and Mouse anti-AP2α (DHSB), applied at dilutions of 1:200, 1:1000, and 1:100 respectively, to label RGCs, jGCaMP8s, and amacrine cells. Secondary antibodies were applied at a dilution of 1:500 and incubated for one hour at room temperature. The retinas were washed again three times in PBS for 30 minutes before flat mount with a coverslip placed on top. Immunostained wholemount images were captured using a Nikon confocal microscope (ECLIPSE Ti2) with a 20x lens and serial filters (BP410-510 for DAPI, BP520-550 for Alexa Fluor 488, BP565-650 for Cy3, and BP650-750 for Alexa Fluor 647). For RGC counting, 20 fields were sampled to encompass the entire retina using a 20x lens, and the percentage of RGC survival was determined by comparing the number of surviving RGCs in injured eyes to those in the contralateral uninjured eyes. For retinal cross-sections, the eyes were dehydrated in a 30% sucrose solution overnight, then embedded in OCT. Serial sections (14 µm) were cut using a Leica cryostat, collected on Superfrost Plus slides, and stored at –80°C until further processing. Cryosections of the eyes were prepared for immunostaining, and images were obtained using a Nikon confocal microscope (ECLIPSE Ti2) equipped with a 40x water immersion lens.

### Optimized adaptive optics two-photon excitation fluorescence microscopy

A schematic of the AO-TPEFM system is shown (Fig. S1). We optimized our previous two-photon excitation fluorescence microscope^34^ to accurately measure and correct the mouse eye aberration with SO. In brief, the system was equipped with a 920 nm fiber laser (FF Ultra 920, TOPTICA) running at 80MHz pulse rate for the fluorescence excitation. The system dispersion is compensated with internal compensation unit to achieve shortest laser pulse duration. The laser was expanded to slightly overfill the aperture of the deformable mirror (DM, Alpao, DM97-15). DM surface was conjugated to a pair of 5-mm galvonmeter scan mirrors (Sunny Technology, SS9215). The galvo X and Y mirrors were conjugated by a 4-f telescope. Then, the galvo mirror was relayed to the electrically tunable lens (ETL, Optotune, EL-16-40-TC-VIS-5D-C). A 4-f system further relayed the ETL to the mouse cornea and compressed the excitation beam to approximately 1.3 mm in diameter so that a typical SO size is larger than the laser beam size.

For two-photon fluorescence imaging, the emitted fluorescence signals were separated by a dichroic mirror (Semrock, FF560-Di01-25×36) into green and red channels. Suitable band-pass and short-pass filters were placed in front of two photomultiplier tubes (Hamamatsu, H7422-40) to select particular fluorescence wavelength and filter out the excitation laser.

For the wavefront sensing, the emitted fluorescence followed the reverse path of the excitation laser. It was descanned by the gavo mirrors and then reflected by the DM. It will be separated by from the femtosecond laser by a dichroic mirror (Semrock, FF705-Di01-25*36) and then relay to the microlens array (SUSS MicroOptics, 18-00197) of a customized Shack-Hartman wavefront sensor (SHWS). The fluorescence will be focused by the microlens array onto an electron-multiplying charge-coupled device (Andor iXon3 888).

### SOHU glaucoma model

A SO induced mouse glaucoma model is utilized here^5,8,37,98^. For experiment group, 2-month-old mice were anesthetized by an intraperiotoneal (i.p) injection of Avertin (0.3mg/g weight). A tunnel was made by a 32G needle through the cornea of the left eye close to the limbus to reach the anterior chamber without injuring lens or iris. Silicone oil (BAUSCH+LOMB OXANE 1300 Silicone Oil) was injected into the anterior chamber slowly using a homemade sterile glass micropipette until the droplet expanded to a diameter of 1.8 mm to 2.2 mm. Veterinary antibiotic ointment (NEOMYCIN CREAM 0.5% - Christo) was applied to the surface of both left and right eye. For control group, mice received same procedures except for the injection of 2uL of PBS to the anterior chamber instead of SO. During the whole procedure, normal saline were applied to keep the cornea moist.

### IOP measurement

Intraocular pressure (IOP) is measured on the injected eye with a tonometer (iCare TA01i) according to the product instructions every time before *in vivo* imaging. Briefly, mice were first anesthetized with a sustained flow of isoflurane (2% at 2L/minute mixed with oxygen) in a sealed chamber. After the initial anesthesia, mice were taken out from chamber and put on a stage with nose cone delivering isoflurane with lower concentration (1% at 0.4L/minute). Tropicamide ophthalmic solution (Mydrin-P) was applied to the cornea three times to fully dilate the pupils. We noticed that IOP usually doesn’t increase immediately after pupils are fully dilated. Since we will perform *in vivo* experiments later, to avoid the risk of too much anesthesia we put the mice back on a heating pad for recovery from anesthesia after we observed that the pupil is fully dilated. Mice usually waked up after 10 minutes rest and were then anesthetized by i.p injection of xylazine (10 mg/kg) and ketamine (80 mg/kg). Before IOP measurement, clean the surface of cornea with PBS and absorb any remaining liquid on cornea. The average of six measurements by tonometer was considered as one machine-generated reading and three machine-generated reading were recorded for each eye. The mean value of three readings was calculated as the IOP value for this time point. Sometimes the increase of IOP appears later than 10 minutes. We usually won’t wait for too long (no more than 20 minutes after pupil is fully dilated) in order to save time for *in vivo* imaging since the mice gradually developed cataract after anesthesia^99^ which cause great troubles for imaging with high imaging quality. After IOP measurement, mice were ready for *in vivo* imaging.

### Longitudinal *in vivo* imaging of mouse retina with AO correction

After being anesthetized with ketamine and xylazine, mice were mounted on an angle adjuster (MAG-2, NARISHIGE, Japan) through a customized nose mask which comprises of a tooth biting bar for fixing head and a tiny air pipe for delivering isoflurane during *in vivo* imaging. 100ul Texas Red (70KDa-Dextran, 2mg/100ul) is retro-orbitally injected into the mice^48,49^ from the contralateral eye (without saline/SO injection) to label the blood vessel patterns for relocating the same sites on the retina at every imaging session. A small droplet of lubricant eye gel (GenTeal Tears) was applied on the cornea of both eyes and a self-designed 0-diopter rigid contact lens^35^ (Advanced Vision Technologies) was placed on the imaging eye to slow down the cataract. Then the angle adjuster is mounted on a 3-axis translation stage (x,y and z direction). The x-axis and y-axis translation is for aligning the center of SO with center of laser beam, and the z direction is for adjusting the conjugation plane with cornea to achieve optimal AO correction FOV. Mice were kept on a homoeohermic blanket to maintain body temperature (37℃) during imaging.

*In vivo* imaging site is chosen from ventral retina and as far from ONH as possible and we find ∼800 um is the limit of our imaging system in order to achieve effective AO correction, which, by our definition, lies in the mid-peripheral retina. Further distance from ONH will cause the block of laser beam due to large laser input angle which is indicated by the spots pattern on WFS (Fig. S21).

To achieve effective AO correction, the center of laser beam should be carefully aligned with the center of SO to avoid the error caused by SO boundary. There are two steps to achieve accurate alignment of the laser beam with SO. The first step is to roughly align the laser beam with the center of pupil with an infrared conversion viewer (ABRM-2000-x, ADOS-TECH) under a low laser power (<1mW) before the cornea. Low laser power (<1mW) is important to avoid injury caused by stationary laser beam exposure^80^ as it may take some time to align the laser with infrared viewer. The second step is to accurately align with the help of SHWS. After the rough alignment, the blurred structures of large retinal vessels will be visualized in a FOV around 400 *μm* under a laser power of 7mW. Adjust the angle adjuster so that a large retinal vessel structure is moved to the center of FOV. A small FOV of 10 μm on the large vessel is scanned as the guiding star for wavefront measurement. If the SO is not well aligned with the laser, the spots pattern on SHWS will display a curved dark band which is due to the large aberrations of the SO boundary (Fig. 1b) and it is easily imaged the position of SO center based on the curvature of dark band. Adjust the x-axis and y-axis translation stages so that the center of SO is well placed in the center of SHWS. Since the diameter of the laser beam is optimized to be smaller than SO diameter, a good alignment can totally avoid the effect of SO boundary and an intact spots pattern can be seen on the SHWS (Fig. 1b).

After alignment of SO and laser beam, AO correction can be performed to improve the imaging resolution as previous work^34^. Briefly, we measured the wavefront of the descanned Texas Red fluorescence signal from a FOV of 10 μm of a large retinal vessel. The laser power for wavefront measurements is less than 7 mW and the integral time is less than 1 s to avoid any injury caused by laser. Several iterations may be needed to achieve optimal imaging quality.

After the correction, the sharp boundary of large vessels and clear structures of tiny capillaries are visualized in a FOV of 200 *μm*. With the help of clear vascular pattern, the same site can be firstly recorded before saline/SO injection and relocated at 0.5wpi, 1.5wpi, 4.5wpi and 7.5wpi. Usually, 2-3 different sites on the ventral retina are imaged for each mouse. The laser power for imaging is 13 mW with a 200 *μm* FOV at 4 frames per second. 4-8 frames were averaged for one slice based on signal intensity and a stack is comprised of 30-40 slice starting from the surface of inner retina with 3.6 *μm* step.

### RBCs flow speed and flux measurement

To measure the blood flow speed and flux, a commonly approach is used through a line scanning along the center line of a capillary^54^ at 1-2 KHz based on the flow speed of the capillary for 1 s. To avoid laser injury, imaging power is usually set to 7 mW and no more than 13 mW unless the signal level of capillaries is quite low. For each imaging site, 5 capillaries were randomly picked in DP and IP. Less than 5 capillaries were usually measured in SP since the vessel pattern is sparser in the SP than DP and IP. Then the blood flow speed is calculated from the time-spatial imaging of line scanning using the Radon transform method^100^ with a 50 −100 ms window size. For some cases when RBCs are rather sparse, window size should be manually adjusted to avoid calculated wrong flow speed from background plasma. The instantaneous flow speed values in 1s are averaged as the mean flow speed value for a capillary. Flux is manually measured by counting the number of RGCs in the time-spatial images. Relative flow speed is calculated by normalizing the longitudinal flow speed of each capillary to the flow speed at the first time point. Relative flux is calculated by normalizing the longitudinal flux to the mean value of all flux at first time point. Relative flux is defined in this way since the minimum value of flux can be or close to zero for some capillaries at the first time point. Dividing the flux at other time points with a small initial flux results in several huge numbers causing significant bias to the statistical mean value.

### Microglia ramification index and motility index

To quantitatively describe the morphological changes of microglia during the glaucoma, we measured the ramification index (RI) as previous work^62,101^. Specifically, an imaging stack of one retinal site from Cx3Cr1-GFP mice is taken. The slices of the interested microglia from a stack are firstly projected through maximum intensity projection (MIP) and then a threshold is applied to the MIP image to acquire a binary mask. The perimeter and area of the binary mask is measured, and the ramification index is calculated by:

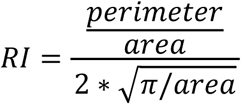

To describe the function of microglia, we use motility index (MI) similar to previous work^61,63,102^. Specifically, 3-5 imaging stacks of one retinal site one site from Cx3Cr1-GFP mice are taken in 20-25 minutes. The MIP images of one microglia of interests at different times are concatenated and registered to create a video. The tip of each microglia process is carefully tracked manually for all the frames with ImageJ Manual Tracking plugins. The instantaneous movement speed of one process is defined as:

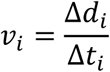

where Δ*d*_*i*_is the processes tips displacement between frame *i* and *i* + 1, and Δ*t*_*i*_ is the time interval between frame *i* and *i* + 1. All the instantaneous movement speed from total video is averaged to get:

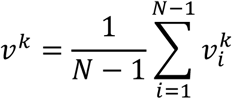

where *k* denotes the k-th process tip. The motility index is defined as the mean value of all tips’ averaged velocity from one microglia:

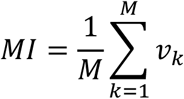

where *M* denotes the total number of process tips of one microglia.

### *In vivo* two-photon RGCs calcium imaging and light stimulation

For RGCs stimulation, a 405nm laser diode is delivered to the mouse eye and form a Maxiwellian scheme^103^ as shown in Fig. S1a. To avoid the contamination of recorded calcium signal due to the stimulation laser^73^, the stimulation laser is synchronized and turned on during the flyback time of imaging scanner with average intensity of 1 *μW*. The frame rate for the calcium imaging is set to 4 Hz and the imaging field of view is 160 *μm*\*160 *μm* on the retina. Stimulation is given after a 10 s delay from the start of two-photon imaging to eliminate the RGCs response (Fig. S15c) to the 920nm femtosecond laser^74^ and the stimulation duration last for 20 s. The total time for one Ca^2+^ imaging session lasts for 60 s. 3 imaging sites in ventral retina are randomly chosen for each mouse. When a Ca^2+^ imaging session is finished on one site, we will wait for at least 3 minutes of dark break before next session on another site to allow full recovery of light-evoked RGC activity^5^.

### Calcium signal analysis

The 60 s Ca^2+^ imaging video is first registered through self-written algorithm in MATLAB. Then the video is average-intensity projected to enhance signal-to-noise ratio and then the mask of each RGCs soma is drawn manually in ImageJ. The time-dependent fluorescent signal from each RGC (*F*) was subtracted with the mean value of fluorescence signal during 10 s pre-stimulus window ( *F*_0_), which can be expressed as Δ*F* = *F* − *F*_0_. The normalized responses ( Δ*F*/*F*_0_) are then denoised by a low-pass filter below normalized passband frequency ( 0.3 *π rad*/*sample*) as previous reports^5^ and used for further analysis.

### RGCs classification

To simply classify RGCs based on the functions, we first determine the range of non-specific Ca^2+^ responses caused by spontaneous activity or imaging procedures but unrelated to UV stimulation similar to previous work^5^. A 60 s video of Ca^2+^ responses from RGCs was recorded without giving UV stimulation. As there was still fluctuation of GCaMP8s fluorescence during the 60 s recording, we separate RGCs into 2 groups based on whether there is increase or decrease of averaged fluorescence intensity during 10-30 s versus that of 0-10 s. The response waveform of all RGCs is averaged separately in each group producing 2 different waveforms with a slight increase or decrease during 60 s period. The upper bound and lower bound of the range of non-specific responses are measured as the maximum value or minimum value of 2 waveforms respectively (Fig. S15b). With UV stimulation, we defined an RGC as no response (NR) if the maximum/minimum value of response is in between the range. For the RGCs beyond this range, we classify them into simple two types of “on-like” and “off-like” by comparing the mean value of Δ*F*/*F*_0_ during 10 - 30 s (stimulation period) with the mean value Δ*F*/*F*_0_ during 5 - 10s (*F*_0_). A RGC is defined as “on-like” if the mean value at stimulation period is larger than *F*_0_, and as “off-like” if the mean value at stimulation period is smaller than *F*_0_.

### Image processing and analysis

The images were processed using MATLAB or ImageJ^104^. To eliminate inter-frame motion artifacts, the images were registered using the StackReg plugin in ImageJ, as well as the self-written algorithm in MATLAB.

The raw data of a volume stack is comprised as a 4-dimensinoal hyperstack (XYTZ). To register the motion artifacts, images on each depth are first registered and averaged to generate a 3-dimensional stack (XYZ), then the 3D stack is registered with adjacent frames at two depths in MATLAB. To register several 3D stacks at different time points for microglia dynamic studies, all the 3D stacks are first merged and then converted to hyperstacks and aligned using GFP signal of microglia with the “Correct 3D drift” plugin in ImageJ. For multi-color channels images, registration is usually based on the red channel (vessel). The transformation matrix is calculated from the red channel and then applied on the images of another channel in MATLAB.

Raw data of RGCs Ca^2+^ slices in the 60 s video are first registered slice by slice with self-written algorithm in MATLAB. The registered video frames were then projected as average intensity projection (AIP). All the images of RGCs in the figures are shown with AIP.

The image pairs with system and full AO correction were shown and processed using the same parameters. For the “system AO” images where the signal intensity was too weak, a linear digital enhancement as indicated in the figures was applied for better visualization.

### Statistical analysis

Statistical analysis and data visualization were performed using GraphPad Prism 10 software and MATLAB. The data are presented as mean ± SD, or ± SEM and α = 0.05 for all analyses. Data normality was first checked using the Shapiro–Wilk normality test. Normally distributed data were analyzed using paired, unpaired two-tailed t-test, one-way and two-way ANOVA test. Non-normally distributed data were analyzed using paired or unpaired non-parametric test (Wilcoxon test, Mixed-effects model analysis with Tukey’s multiple comparisons test). Some data were presented as contingency table. No statistical methods were used to determine the sample size.

## Acknowledgement

The Hong Kong Research Grants Council through grants (16102122, 16102123, 16102421, 16102518, 16102920, T13-607/12R, T13-605/18W, C600217GF, C6001-19E, C6034-21G, T13-602/21N), the Innovation and Technology Commission (ITCPD/17-9), the Area of Excellence Scheme of the University Grants Committee (AoE/M-604/16, AOE/M-09/12) and the Hong Kong University of Science & Technology (HKUST) through grant 30 for 30 Research Initiative Scheme. We thank the Hong Kong Center for Neurodegenerative Diseases InnoHK of Hong Kong SAR and the histology and microscopy core of Biosciences Central Research Facility, HKUST (Clear Water Bay).

## Author contributions

Y.F., S.H. P.B.M., T. X. and J.Y.Q. conceived the research idea and designed the experiments; Y.F., Z.Q. built and optimized the imaging systems; Y.F. performed animal surgery and imaging experiments; P.B.M. performed the Brn3b-Cre line creation, SO injection, virus injection and histology study under the supervision of T.X.; Y.F. and P.B.M. analyzed the data with the help of Y.H.; Y.F. and P.B.M. wrote the paper with inputs from all other authors.

## References

1. Weinreb, R. N., Aung, T. & Medeiros, F. A. The pathophysiology and treatment of glaucoma: A review. JAMA 311, 1901–1911 (2014).

2. Tham, Y. C. et al. Global prevalence of glaucoma and projections of glaucoma burden through 2040: A systematic review and meta-analysis. Ophthalmology 121, 2081–2090 (2014).

3. Jayaram, H., Kolko, M., Friedman, D. S. & Gazzard, G. Glaucoma: now and beyond. The Lancet vol. 402 1788–1801 Preprint at 10.1016/S0140-6736(23)01289-8 (2023).

4. Singh, K. & Shrivastava, A. Intraocular pressure fluctuations: how much do they matter? Curr Opin Ophthalmol 20, 84–87 (2009).

5. Li, L. et al. Longitudinal in vivo Ca 2+ imaging reveals dynamic activity changes of diseased retinal ganglion cells at the single-cell level. Proceedings of the National Academy of Sciences 119, 2017 (2022).

6. Casson, R. J., Chidlow, G., Wood, J. P. M., Crowston, J. G. & Goldberg, I. Definition of glaucoma: Clinical and experimental concepts. Clin Exp Ophthalmol 40, 341–349 (2012).

7. Quigley, H. A. Neuronal death in glaucoma. Prog Retin Eye Res 18, 39–57 (1999).

8. Zhang, J. et al. Silicone oil-induced ocular hypertension and glaucomatous neurodegeneration in mouse. Elife 8, 1–19 (2019).

9. Mukai, R. et al. Mouse model of ocular hypertension with retinal ganglion cell degeneration. PLoS One 14, 1–21 (2019).

10. Flammer, J. et al. The impact of ocular blood flow in glaucoma. Prog Retin Eye Res 21, 359–393 (2002).

11. Burgansky–Eliash, Z., Bartov, E., Barak, A., Grinvald, A. & Gaton, D. Blood-Flow Velocity in Glaucoma Patients Measured with the Retinal Function Imager. Curr Eye Res 41, 965–970 (2016).

12. Alarcon-Martinez, L. et al. Pericyte dysfunction and loss of interpericyte tunneling nanotubes promote neurovascular deficits in glaucoma. Proc Natl Acad Sci U S A 119, 1–11 (2022).

13. Jeon, S. J., Shin, D. Y., Park, H. Y. L. & Park, C. K. Association of Retinal Blood Flow with Progression of Visual Field in Glaucoma. Sci Rep 9, 1–8 (2019).

14. Tanito, M., Kaidzu, S., Takai, Y. & Ohira, A. Association between systemic oxidative stress and visual field damage in open-angle glaucoma. Sci Rep 6, (2016).

15. Izzotti, A., Bagnis, A. & Saccà, S. C. The role of oxidative stress in glaucoma. Mutation Research - Reviews in Mutation Research vol. 612 105–114 Preprint at 10.1016/j.mrrev.2005.11.001 (2006).

16. Tanito, M., Kaidzu, S., Takai, Y. & Ohira, A. Association between systemic oxidative stress and visual field damage in open-angle glaucoma. Sci Rep 6, (2016).

17. Wiggs, J. L. & Pasquale, L. R. Genetics of glaucoma. Human Molecular Genetics vol. 26 R21–R27 Preprint at 10.1093/hmg/ddx184 (2017).

18. Jiang, S., Kametani, M. & Chen, D. F. Adaptive Immunity: New Aspects of Pathogenesis Underlying Neurodegeneration in Glaucoma and Optic Neuropathy. Frontiers in Immunology vol. 11 Preprint at 10.3389/fimmu.2020.00065 (2020).

19. The role of microglia in the progression of glaucomatous neurodegeneration-a review. Int J Ophthalmol (2018) doi:10.18240/ijo.2018.01.22.

20. Zhao, X., Sun, R., Luo, X., Wang, F. & Sun, X. The Interaction Between Microglia and Macroglia in Glaucoma. Front Neurosci 15, (2021).

21. Zhi, Z., Cepurna, W. O., Johnson, E. C., Morrison, J. C. & Wang, R. K. Impact of intraocular pressure on changes of blood flow in the retina, choroid, and optic nerve head in rats investigated by optical microangiography. Biomed Opt Express 3, 2220 (2012).

22. Emre, M. Ocular blood flow alteration in glaucoma is related to systemic vascular dysregulation. British Journal of Ophthalmology 88, 662–666 (2004).

23. Nicolela, M. T., Hnik, P. & Drance, S. M. Scanning Laser Doppler Flowmeter Study of Retinal and Optic Disk Blood Flow in Glaucomatous Patients. Am J Ophthalmol 122, 775–783 (1996).

24. Rao, H. L. et al. Optical Coherence Tomography Angiography in Glaucoma. J Glaucoma 29, 312–321 (2020).

25. Alarcon-Martinez, L. et al. Interpericyte tunnelling nanotubes regulate neurovascular coupling. Nature 585, 91–95 (2020).

26. Xu, H., Chen, M., Forrester, J. V. & Lois, N. Cataract Surgery Induces Retinal Pro-inflammatory Gene Expression and Protein Secretion. Investigative Opthalmology & Visual Science 52, 249 (2011).

27. Wei, X., Cho, K. S., Thee, E. F., Jager, M. J. & Chen, D. F. Neuroinflammation and microglia in glaucoma: time for a paradigm shift. J Neurosci Res 97, 70–76 (2019).

28. Ramírez, A. I. et al. Time course of bilateral microglial activation in a mouse model of laser-induced glaucoma. Sci Rep 10, 1–17 (2020).

29. Geng, Y. et al. Optical properties of the mouse eye. Biomed Opt Express 2, 717 (2011).

30. Ou, Y., Jo, R. E., Ullian, E. M., Wong, R. O. L. & Della Santina, L. Selective vulnerability of specific retinal ganglion cell types and synapses after transient ocular hypertension. Journal of Neuroscience 36, 9240–9252 (2016).

31. Santina, L. Della, Inman, D. M., Lupien, C. B., Horner, P. J. & Wong, R. O. L. Differential progression of structural and functional alterations in distinct retinal ganglion cell types in a mouse model of glaucoma. Journal of Neuroscience 33, 17444–17457 (2013).

32. Helmchen, F. & Denk, W. Deep tissue two-photon microscopy. Nat Methods 2, 932–940 (2005).

33. Svoboda, K. & Yasuda, R. Principles of Two-Photon Excitation Microscopy and Its Applications to Neuroscience. Neuron vol. 50 823–839 Preprint at 10.1016/j.neuron.2006.05.019 (2006).

34. Qin, Z. et al. Adaptive optics two-photon microscopy enables near-diffraction-limited and functional retinal imaging in vivo. Light Sci Appl 9, 79 (2020).

35. Zhang, Q. et al. Retinal microvascular and neuronal pathologies probed in vivo by adaptive optical two-photon fluorescence microscopy. Elife 12, 1–23 (2023).

36. Fang, F. et al. Chronic mild and acute severe glaucomatous neurodegeneration derived from silicone oil-induced ocular hypertension. Sci Rep 11, 9052 (2021).

37. Zhang, J. et al. A Reversible Silicon Oil-Induced Ocular Hypertension Model in Mice. Journal of Visualized Experiments (2019) doi:10.3791/60409.

38. Shin, J. Y. et al. HIGHER-ORDER ABERRATIONS IN EYES WITH SILICONE OIL TAMPONADE. Retina 40, 735–742 (2020).

39. Fruttiger, M. Development of the retinal vasculature. Angiogenesis 10, 77–88 (2007).

40. Simmons, A. B. & Fuerst, P. G. Analysis of Retinal Vascular Plexuses and Interplexus Connections. in 317–330 (2018). doi:10.1007/978-1-4939-7720-8_22.

41. Stefánsson, E., Anderson, M. M., Landers, M. B., Tiedeman, J. S. & McCuen, B. W. REFRACTIVE CHANGES FROM USE OF SILICONE OIL IN VITREOUS SURGERY. Retina 8, 20–23 (1988).

42. Smith, R. C., Smith, G. T. & Wong, D. Refractive changes in silicone filled eyes. Eye 4, 230–234 (1990).

43. So, P. T. C., Dong, C. Y., Masters, B. R. & Berland, K. M. Two-photon excitation fluorescence microscopy. Annu Rev Biomed Eng 2, 399–429 (2000).

44. Tanito, M., Itai, N., Ohira, A. & Chihara, E. Reduction of posterior pole retinal thickness in glaucoma detected using the Retinal Thickness Analyzer. Ophthalmology 111, 265–275 (2004).

45. Greenfield, D. S. Macular Thickness Changes in Glaucomatous Optic Neuropathy Detected Using Optical Coherence Tomography. Archives of Ophthalmology 121, 41 (2003).

46. Leung, C. K. et al. Evaluation of Retinal Nerve Fiber Layer Progression in Glaucoma: A Study on Optical Coherence Tomography Guided Progression Analysis. Investigative Opthalmology & Visual Science 51, 217 (2010).

47. Ferguson, L. R., Dominguez, J. M., Balaiya, S., Grover, S. & Chalam, K. V. Retinal Thickness Normative Data in Wild-Type Mice Using Customized Miniature SD-OCT. PLoS One 8, 1–8 (2013).

48. Yardeni, T., Eckhaus, M., Morris, H. D., Huizing, M. & Hoogstraten-Miller, S. Retro-orbital injections in mice. Lab Anim (NY) 40, 155–160 (2011).

49. Li, S. et al. Retro-orbital injection of FITC-dextran is an effective and economical method for observing mouse retinal vessels. Mol Vis 17, 3566–3573 (2011).

50. Shin, J. W., Sung, K. R., Lee, G. C., Durbin, M. K. & Cheng, D. Ganglion Cell–Inner Plexiform Layer Change Detected by Optical Coherence Tomography Indicates Progression in Advanced Glaucoma. Ophthalmology 124, 1466–1474 (2017).

51. Mahmoudinezhad, G. et al. Comparison of Ganglion Cell Layer and Ganglion Cell/Inner Plexiform Layer Measures for Detection of Early Glaucoma. Ophthalmol Glaucoma 6, 58–67 (2023).

52. Nicolela, M. T., Hnik, P. & Drance, S. M. Scanning Laser Doppler Flowmeter Study of Retinal and Optic Disk Blood Flow in Glaucomatous Patients. Am J Ophthalmol 122, 775–783 (1996).

53. Holló, G., van den Berg, T. J. T. P. & Greve, E. L. Scanning laser Doppler flowmetry in glaucoma. Int Ophthalmol 20, 63–70 (1996).

54. Zhong, Z., Petrig, B. L., Qi, X. & Burns, S. A. In vivo measurement of erythrocyte velocity and retinal blood flow using adaptive optics scanning laser ophthalmoscopy. Opt Express 16, 12746 (2008).

55. Hartmann, D. A. et al. Brain capillary pericytes exert a substantial but slow influence on blood flow. Nat Neurosci 24, 633–645 (2021).

56. Naskar, R., Wissing, M. & Thanos, S. Detection of early neuron degeneration and accompanying microglial responses in the retina of a rat model of glaucoma. Invest Ophthalmol Vis Sci 43, 2962–2968 (2002).

57. Blank, T. et al. Early Microglia Activation Precedes Photoreceptor Degeneration in a Mouse Model of CNGB1-Linked Retinitis Pigmentosa. Front Immunol 8, (2018).

58. Holloway, O. G., Canty, A. J., King, A. E. & Ziebell, J. M. Rod microglia and their role in neurological diseases. Semin Cell Dev Biol 94, 96–103 (2019).

59. Bosco, A. et al. Neurodegeneration severity can be predicted from early microglia alterations monitored in vivo in a mouse model of chronic glaucoma. DMM Disease Models and Mechanisms 8, 443–455 (2015).

60. Avignone, E., Lepleux, M., Angibaud, J. & Nägerl, U. V. Altered morphological dynamics of activated microglia after induction of status epilepticus. J Neuroinflammation 12, (2015).

61. Gyoneva, S. et al. Systemic inflammation regulates microglial responses to tissue damage in vivo. Glia 62, 1345–1360 (2014).

62. Madry, C. et al. Microglial Ramification, Surveillance, and Interleukin-1β Release Are Regulated by the Two-Pore Domain K+ Channel THIK-1. Neuron 97, 299–312.e6 (2018).

63. Nimmerjahn, A., Kirchhoff, F. & Helmchen, F. Resting Microglial Cells Are Highly Dynamic Surveillants of Brain Parenchyma in Vivo. Science (1979) 308, 1314–1318 (2005).

64. Fang, R. et al. Three-dimensional single-cell transcriptome imaging of thick tissues. 12, 90029 (2023).

65. Chen, A. et al. Spatiotemporal transcriptomic atlas of mouse organogenesis using DNA nanoball-patterned arrays. Cell 185, 1777–1792.e21 (2022).

66. Guo, L., Choi, S., Bikkannavar, P. & Cordeiro, M. F. Microglia: Key Players in Retinal Ageing and Neurodegeneration. Front Cell Neurosci 16, (2022).

67. Rojas, B. et al. Microglia in mouse retina contralateral to experimental glaucoma exhibit multiple signs of activation in all retinal layers. J Neuroinflammation 11, 133 (2014).

68. Badea, T. C., Cahill, H., Ecker, J., Hattar, S. & Nathans, J. Distinct Roles of Transcription Factors Brn3a and Brn3b in Controlling the Development, Morphology, and Function of Retinal Ganglion Cells. Neuron 61, 852–864 (2009).

69. Zhang, Y., et al. *Fast and Sensitive GCaMP Calcium Indicators for Imaging Neural Populations*. Nature vol. 615 (Springer US, 2023).

70. Applebury, M. L. et al. The Murine Cone Photoreceptor. Neuron 27, 513–523 (2000).

71. Baden, T. et al. The functional diversity of retinal ganglion cells in the mouse. Nature 529, 345–350 (2016).

72. Nikonov, S. S., Kholodenko, R., Lem, J. & Pugh, E. N. Physiological Features of the S- and M-cone Photoreceptors of Wild-type Mice from Single-cell Recordings. J Gen Physiol 127, 359–374 (2006).

73. Yoshimatsu, T. et al. Ancestral circuits for vertebrate color vision emerge at the first retinal synapse. Sci Adv 7, (2021).

74. Euler, T. et al. Eyecup scope-optical recordings of light stimulus-evoked fluorescence signals in the retina. Pflugers Arch 457, 1393–1414 (2009).

75. Leung, C. K. S. et al. Long-term in vivo imaging and measurement of dendritic shrinkage of retinal ganglion cells. Invest Ophthalmol Vis Sci 52, 1539–1547 (2011).

76. Wang, A. Y. M. et al. Potential mechanisms of retinal ganglion cell type-specific vulnerability in glaucoma. Clin Exp Optom 103, 562–571 (2020).

77. Feng, L. et al. Sustained ocular hypertension induces dendritic degeneration of mouse retinal ganglion cells that depends on cell type and location. Invest Ophthalmol Vis Sci 54, 1106–1117 (2013).

78. Choi, J. et al. Spatial organization of the mouse retina at single cell resolution by MERFISH. Nat Commun 14, 4929 (2023).

79. Wiggs, J. L. Genetic Etiologies of Glaucoma. Archives of Ophthalmology 125, 30 (2007).

80. Jayabalan, G. S. et al. Retinal safety evaluation of two-photon laser scanning in rats. Biomed Opt Express 10, 3217 (2019).

81. Masters, B. R. et al. Mitigating thermal mechanical damage potential during two-photon dermal imaging. J Biomed Opt 9, 1265 (2004).

82. Delori, F. C., Webb, R. H. & Sliney, D. H. Maximum permissible exposures for ocular safety (ANSI 2000), with emphasis on ophthalmic devices. Journal of the Optical Society of America A 24, 1250 (2007).

83. Yang, G., Pan, F., Parkhurst, C. N., Grutzendler, J. & Gan, W. B. Thinned-skull cranial window technique for long-term imaging of the cortex in live mice. Nat Protoc 5, 213–220 (2010).

84. Drew, P. J. et al. Chronic optical access through a polished and reinforced thinned skull. Nat Methods 7, 981–984 (2010).

85. Fernández-Arjona, M. del M., Grondona, J. M., Fernández-Llebrez, P. & López-Ávalos, M. D. Microglial Morphometric Parameters Correlate With the Expression Level of IL-1β, and Allow Identifying Different Activated Morphotypes. Front Cell Neurosci 13, (2019).

86. Soltys, Z., Ziaja, M., Pawlinski, R., Setkowicz, Z. & Janeczko, K. Morphology of reactive microglia in the injured cerebral cortex. Fractal analysis and complementary quantitative methods. J Neurosci Res 63, 90–97 (2001).

87. Zborowski-Gutman, L., Treister, G., Naveh, N., Chen, V. & Blumenthal, M. Acute glaucoma following vitrectomy and silicone oil injection. British Journal of Ophthalmology 71, 903–906 (1987).

88. Joseph, A., Guevara-Torres, A. & Schallek, J. Imaging single-cell blood flow in the smallest to largest vessels in the living retina. Elife 8, 533919 (2019).

89. Anderson, D. R. Glaucoma, Capillaries and Pericytes. Ophthalmologica 210, 257–262 (1996).

90. Hughes, S. et al. Altered pericyte-endothelial relations in the rat retina during aging: Implications for vessel stability. Neurobiol Aging 27, 1838–1847 (2006).

91. Haruwaka, K. et al. Dual microglia effects on blood brain barrier permeability induced by systemic inflammation. Nat Commun 10, 1–17 (2019).

92. Ahmad, I. & Subramani, M. Microglia: Friends or Foes in Glaucoma? A Developmental Perspective. Stem Cells Transl Med 11, 1210–1218 (2022).

93. Masuda, T., Croom, D., Hida, H. & Kirov, S. A. Capillary blood flow around microglial somata determines dynamics of microglial processes in ischemic conditions. Glia 59, 1744–1753 (2011).

94. Eyo, U. & Dailey, M. E. Effects of oxygen-glucose deprivation on microglial mobility and viability in developing mouse hippocampal tissues. Glia 60, 1747–1760 (2012).

95. Sanes, J. R. & Masland, R. H. The Types of Retinal Ganglion Cells: Current Status and Implications for Neuronal Classification. Annu Rev Neurosci 38, 221–246 (2015).

96. Tien, N.-W., Pearson, J. T., Heller, C. R., Demas, J. & Kerschensteiner, D. Genetically Identified Suppressed-by-Contrast Retinal Ganglion Cells Reliably Signal Self-Generated Visual Stimuli. Journal of Neuroscience 35, 10815–10820 (2015).

97. Murphy, G. J. & Rieke, F. Network Variability Limits Stimulus-Evoked Spike Timing Precision in Retinal Ganglion Cells. Neuron 52, 511–524 (2006).

98. Fang, F. et al. Chronic mild and acute severe glaucomatous neurodegeneration derived from silicone oil-induced ocular hypertension. Sci Rep 11, 9052 (2021).

99. Acute lens opacity induced by different kinds of anesthetic drugs in mice. Int J Ophthalmol 12, (2019).

100. Drew, P. J., Blinder, P., Cauwenberghs, G., Shih, A. Y. & Kleinfeld, D. Rapid determination of particle velocity from space-time images using the Radon transform. J Comput Neurosci 29, 5–11 (2010).

101. Wu, W. et al. In vivo imaging in mouse spinal cord reveals that microglia prevent degeneration of injured axons. Nat Commun 15, 8837 (2024).

102. Joseph, A., Power, D. & Schallek, J. Imaging the dynamics of individual processes of microglia in the living retina in vivo. Biomed Opt Express 12, 6157 (2021).

103. Yin, L. et al. Imaging light responses of retinal ganglion cells in the living mouse eye. J Neurophysiol 109, 2415–2421 (2013).

104. Schindelin, J., et al. Fiji: an open-source platform for biological-image analysis. Nat Methods 9, 676–682 (2012).

